# Identification of compounds that repress DUX4 expression in facioscapulohumeral muscular dystrophy

**DOI:** 10.64898/2026.03.09.710626

**Authors:** Ning Chang, Hannah P. Moore, Charis L. Himeda, Terrence E. O’Brien, William Thomas, Baback Roshanravan, Takako I. Jones, Peter L. Jones

## Abstract

Facioscapulohumeral muscular dystrophy (FSHD) is caused by epigenetic dysregulation of the disease locus, leading to pathogenic misexpression of *DUX4* in skeletal muscle. Thus, most FSHD therapeutic approaches target *DUX4*. Our previous study identified the chromatin remodeling factor BAZ1A (bromodomain adjacent to zinc finger domain protein 1A) as a promising target for therapeutic development. Here we used an artificial intelligence-based screening pipeline to identify molecules predicted to bind the BAZ1A bromodomain, and validated hit compounds using FSHD-specific assays in FSHD myocytes. One compound, termed C06, emerged as a potent and specific repressor of *DUX4* and DUX4 target gene expression. Interestingly, while C06 exhibited binding to BAZ1A in vitro, it can also inhibit multiple kinases, including p38α, an upstream activator of *DUX4*. Despite this, at low doses C06 was an equally effective and more specific repressor of *DUX4* than losmapimod, which is a robust and specific p38 inhibitor. Thus, C06 is a useful tool for potent and specific *DUX4* suppression, and a viable candidate for further development. Our results highlight both the utility and limitations of AI for targeted drug discovery, and the importance of using an FSHD-specific functional screening strategy for selecting relevant candidates.

## Introduction

Facioscapulohumeral muscular dystrophy (FSHD, OMIM: 158900, 158901) is the third most common muscular dystrophy, and one for which no cures or treatments exist^1,2^. All forms of the disease are caused by epigenetic dysregulation of the D4Z4 macrosatellite repeat array at chromosome 4q35, which is normally silenced in adult somatic cells^3,4^. FSHD1, the most common form of the disease, is linked to contractions at this array^5–7^, resulting in chromatin relaxation. FSHD2 is caused by mutations in proteins that maintain epigenetic silencing of the D4Z4 array, leading to a similar chromatin relaxation^8–10^. In both forms of the disease, loss of epigenetic repression leads to aberrant expression of the Double Homeobox 4 (*DUX4)* retrogene in skeletal muscle^11^. DUX4, a cleavage-stage pioneer transcription factor, activates an early embryonic program, which causes pathology when misexpressed in adult skeletal muscle^4,12,13^. Although an intact *DUX4* open reading frame resides in every D4Z4 repeat unit in the macrosatellite array^14^, only the full-length *DUX4* mRNA *(DUX4-fl)* encoded by the distal-most repeat is stably expressed and translated due to the polyadenylation signal (PAS) residing in an exon distal to the array in disease-permissive alleles^11,15^.

As a dominant gain-of-function disease, FSHD is amenable to many viable *DUX4*-targeting therapeutic strategies, which include CRISPR-based approaches, antisense technologies, and small molecule inhibitors^16^. Targeting the *DUX4* PAS using CRISPR/Cas9 gene editing is extremely inefficient^17–19^ and carries the attendant risk of off-target cutting. While different CRISPR inhibition strategies are under development^20^ and one is currently being tested in a Phase1/2 clinical trial (NCT06907875), CRISPR-based muscle gene therapies face a number of serious challenges, including the need to achieve efficient delivery and prevent Cas immunity. Many investigators have focused on developing *DUX4* antisense approaches in preclinical studies^21^ and in two ongoing trials (NCT05747924; NCT06131983); these are promising, but still face the challenge of achieving both safety and efficacy over a lifetime of chronic administration.

While small molecules remain the gold standard for treatment of most disorders, the unique aspects of FSHD create complications not present in traditional drug screening^16^. FSHD muscle cells typically exhibit very low and sporadic expression of *DUX4-fl* mRNA and DUX4-FL protein; detecting a decrease in levels of a low-abundance transcription factor presents difficulties for traditional high-throughput screening (HTS) of chemical libraries. Thus, many investigators identify lead compounds through indirect expression screens using DUX4 reporter constructs, as opposed to directly measuring *DUX4-fl* transcript levels. This screening method is limited by the content of chemical libraries, dosing, and modes of action. Despite several promising proof-of-concept studies^16^, only one small molecule has been in clinical trials for FSHD: the p38 inhibitor losmapimod^22,23^, a repurposed cardiovascular drug which failed its primary endpoint in Phase 3 (NCT04003974) and has always raised concerns regarding the detrimental effects of chronically targeting a key regulator of muscle biology in dystrophic muscles^24,25^. Thus, it is clear that better and safer targets for small molecule development in FSHD are needed.

In contrast to a chemical library screen, Himeda *et al.* performed a candidate-based screen in FSHD myocytes to identify potential targets for informed drug design^26^. Candidates were chosen based on their known roles in regulating active chromatin states, and narrowed to those that: 1) are not global regulators of transcription, and 2) possess domains that are likely to be druggable. Knockdown of several of these epigenetic regulators by either short hairpin RNAs or CRISPR inhibition increased chromatin repression at the D4Z4 array and successfully repressed expression of *DUX4-fl* and DUX4-FL target genes^26^, suggesting that these factors may be viable candidates for FSHD therapeutic drug development. SMARCA5 (SWI/SNF related, matrix associated, actin dependent regulator of chromatin, subfamily A, member 5; SNF2H) and BAZ1A (bromodomain adjacent to zinc finger domain 1A; ACF1)^26^ emerged as two of the top candidates from this screen.

Interestingly, SMARCA5 and BAZ1A are both components of the ACF complex, which is a member of the ISWI (imitation switch) family of ATP-dependent chromatin remodeling complexes^27^. Although non-enzymatic, BAZ1A plays a key regulatory role within this complex, controlling the directionality, efficiency, and specificity of the SMARCA5 ATPase, which utilizes ATP hydrolysis to remodel nucleosomes, controlling access to chromatin^28,29^. While both factors contain druggable domains, targeting the ATPase domain of SMARCA5 carries a higher risk of off-target effects, as the ATPase domain is a common and essential feature across numerous proteins involved in diverse cellular processes. By contrast, the bromodomain of BAZ1A is a structural motif only found in certain chromatin-associated factors, where it functions as a reader of acetylated lysine residues on histones and other proteins. Moreover, transgenic mice harboring conditional knockout alleles of *BAZ1A* revealed that haploinsufficiency is generally well-tolerated (PL Jones lab, manuscript in preparation), suggesting that inhibition of BAZ1A represents a viable therapeutic strategy. In this study, we used an artificial intelligence (AI)-based approach to identify small molecules predicted to interact with the BAZ1A bromodomain, and validated these compounds in FSHD-specific assays.

## Results

To identify potential small molecule inhibitors of BAZ1A, we collaborated with Atomwise, Inc., a company specializing in AI-driven drug discovery. Using deep learning algorithms, the AtomNet^®^ platform developed by Atomwise, Inc. predicts the binding affinities of small molecules to protein targets^30^. Using this AI model, we virtually screened approximately one million chemical structures for their predicted ability to block the BAZ1A bromodomain, resulting in 72 top-scoring compounds (Figure 1A). To validate these compounds, we performed an FSHD-specific screen using an immortalized patient muscle cell line (15Ai) that expresses *DUX4-fl* from the endogenous locus. Since *DUX4-fl* is only expressed in differentiated myocytes^31^, myoblasts must first undergo myogenic differentiation to allow *DUX4-fl* expression prior to analysis. Given the variability in differentiation kinetics among patient cell lines, we determined the differentiation time course for the 15Ai line to identify the window of peak *DUX4-fl* expression within the first four days of differentiation, when the cells are still healthy in culture (Figure 1B). As expected, *DUX4-fl* was almost undetectable in undifferentiated myoblasts, whereas switching to differentiation conditions significantly induced *DUX4-fl* expression in a time-dependent manner (Figure 1B). The time course analysis indicated that 4 days of differentiation provided an optimal window for the drug screening assay (Figure 1C).

**Figure 1.**
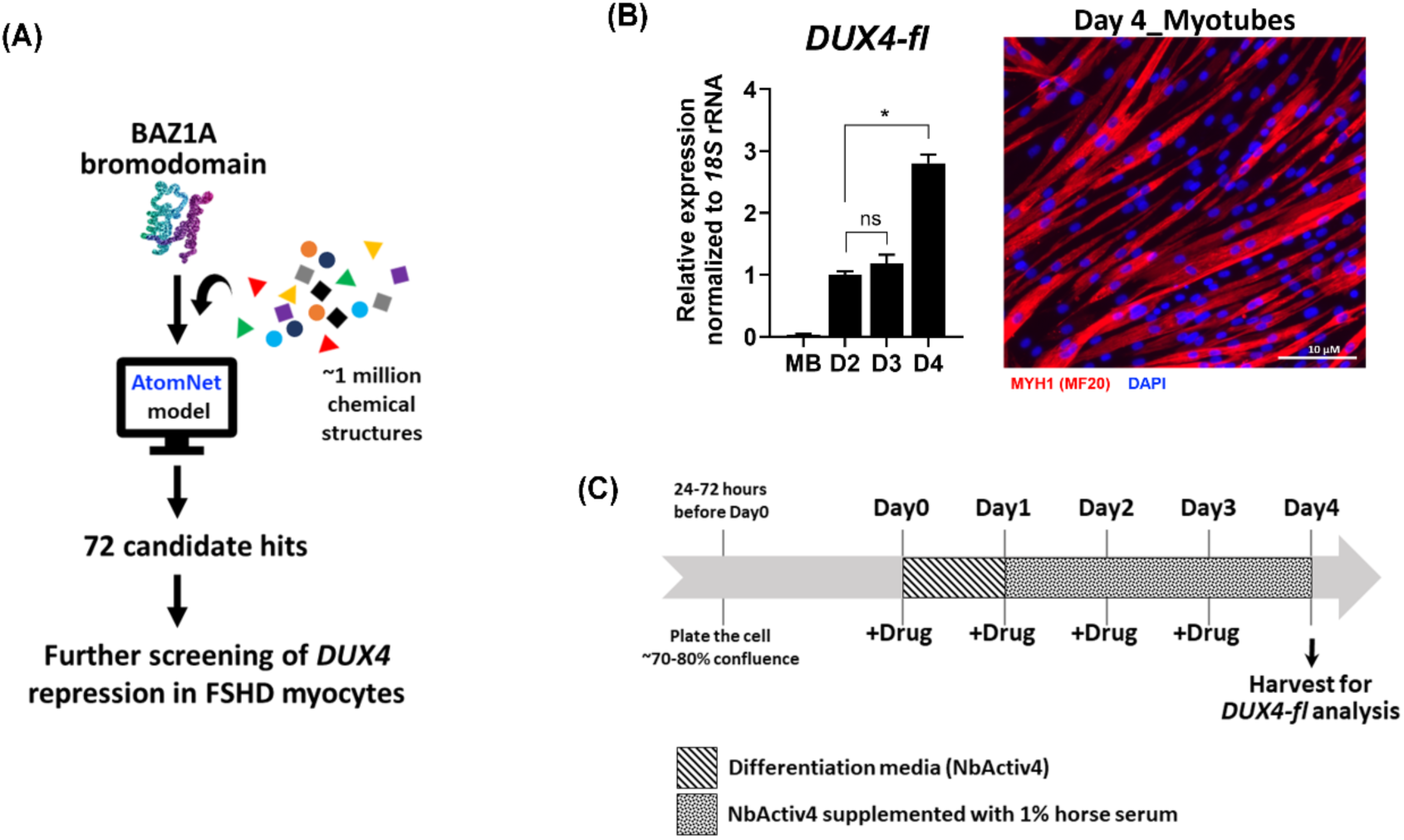
Screening for BAZ1A small-molecule inhibitors. (**A**) Experimental pipeline for identification of BAZ1A small-molecule inhibitors. The AtomNet^®^ (AtomWise, Inc) AI model was used to search ∼1 million commercially available chemical structures for predicted binding to the BAZ1A bromodomain. The top 72 hit compounds were tested for their ability to repress *DUX4* expression in FSHD myocytes. **(B)** Timecourse expression of *DUX4* full-length (*DUX4-fl*) mRNA was assayed by RT-qPCR in FSHD1 myocytes (15Ai). Proliferating myoblasts (MB) were differentiated into myotubes over the course of 4 days. Expression levels of *DUX4-fl* at each time point were normalized to *18S* rRNA. Data are plotted as the mean ± standard error of the mean (SEM) of three independent experiments, with average expression at Day2 (D2) set to 1. Representative image of myotubes at day 4 of differentiation was visualized using immunofluorescence staining of human myosin heavy chain 1 (MYH1). Scale bar = 10 µM. **(C)** The FSHD-customized protocol for validating candidates identified in the AI-based screen.

We next evaluated whether the compounds predicted to inhibit BAZ1A could suppress *DUX4-fl* expression. 15Ai myoblasts were induced to differentiate and treated with each compound at 1 μM for four days. The p38 inhibitor losmapimod was included as a positive control for *DUX4-fl* repression^22^, and its efficacy was confirmed across multiple FSHD cell lines using our protocol (Supplementary Figure S1). All compounds received from Atomwise, Inc. were coded for a blinded screen. Eight compounds (A01, B10, C01, C03, C06, D07, D08, D09) exhibited >50% repression of *DUX4-fl* expression, with C06 exhibiting the most robust repression (Figure 2A). The chemical structures of these compounds are shown in Figure 2B and Table 1. As levels of DUX4-FL protein are low and difficult to assess in FSHD myocytes, we routinely assess levels of DUX4-FL target gene expression as the more reliable assay and relevant functional readout of DUX4-FL activity. Surprisingly, only 3 of the 8 compounds (A01, C06, D08) effectively attenuated the expression of two DUX4-FL target genes, *MBD3L2* and *TRIM43,* with C06 exhibiting by far the most robust repression (Figure 2C). Importantly, all 8 compounds at 1 μM displayed minimal or no effect on myosin heavy chain 1 (*MYH1*) expression (Figure 2D), indicating that the observed *DUX4-fl* reduction was not an indirect effect of impaired myogenesis. By contrast, losmapimod led to a striking decrease in *MYH1* levels (Figure 2D), which is unsurprising, given the key role of p38 in muscle differentiation^24,25^. When we compared the fusion index for FSHD1 muscle cells treated with C06 or losmapimod, we found that at the high concentration (1 µM), losmapimod greatly impaired muscle fusion (∼80% reduction), whereas C06 produced only a slight effect (∼25% reduction) (Supplementary Figure S2A). Even at the lower concentration (100 nM), losmapimod still inhibited muscle fusion, whereas C06 showed no effect (Supplementary Figure S2A). Similar results were observed in a healthy primary muscle cell line (Supplementary Figure S2B). Due to its specificity and extremely potent ability to repress *DUX4-fl* and its downstream targets, we chose to move forward with C06.

**Figure 2.**
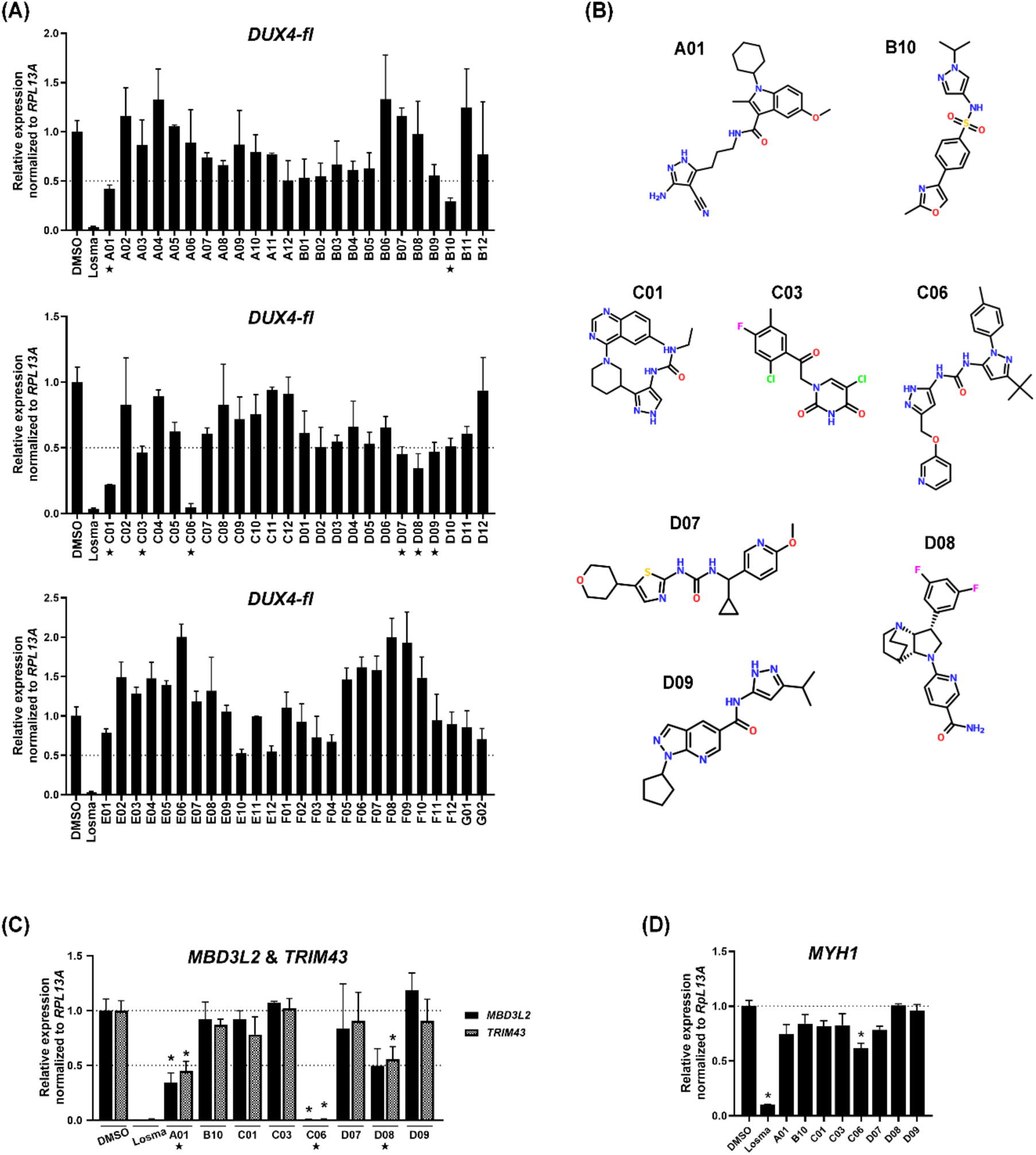
Validation of compounds suppressing *DUX4-fl* in FSHD1 myocytes. (**A, C and D)** FSHD1 myoblasts (15Ai) were differentiated and treated at 1 µM with individual compounds from the AI screen (A01 to G02) or losmapimod (Losma) as a positive control for *DUX4* repression. Expression levels of *DUX4-fl* (A), DUX4-FL target genes *MBD3L2* and *TRIM43* (C), and the differentiation marker *MYH1* (D) were assessed by RT-qPCR and normalized to *RPL13A*. Data are plotted as the mean ± standard error of the mean (SEM) of two or three independent experiments, with average expression of vehicle-treated cells (DMSO) set to 1. \**p* < 0.05 is from comparing to DMSO. **(B)** Chemical structures of compounds (asterisks in A) that yielded at least 50% repression (dashed lines in A) of *DUX4-fl*.

**Table 1.**
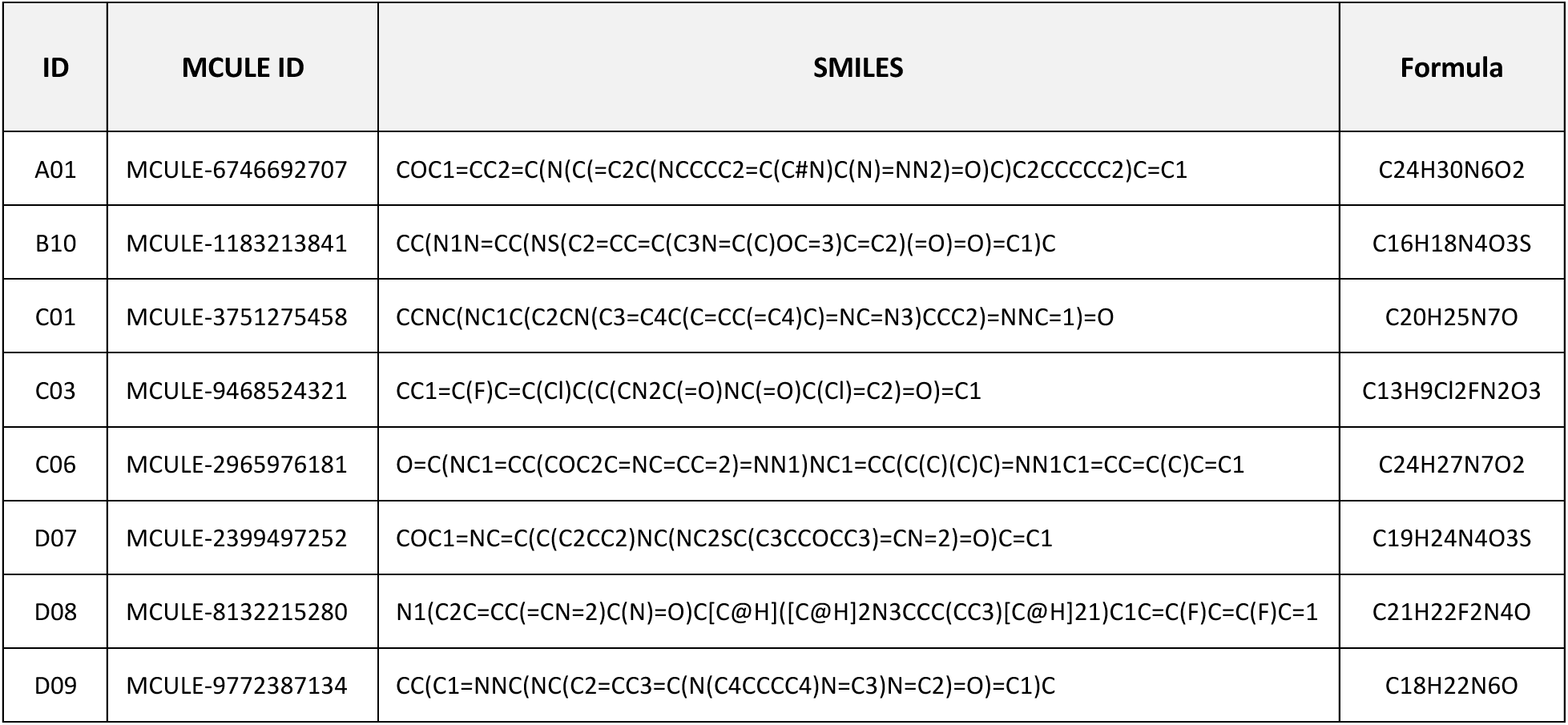
Chemical structures of top 8 hit compounds predicted to bind the BAZ1A bromodomain.

We next investigated the dose dependence of C06-mediated *DUX4-fl* repression. C06 exhibited a clear dose-dependent repression of *DUX4-fl* and its downstream targets, with IC50s of 23.78 nM for *DUX4*, 22.18 nM for *MBD3L2*, and 21.89 nM for *TRIM43* (Figure 3). Importantly, concentrations of C06 that effectively suppressed *DUX4-fl* did not adversely affect *MYH1* expression, which showed an IC50 of 81 nM, indicating that the effect of C06 on *DUX4-fl* is not due to impaired myogenesis (Figure 3). To determine whether the effects of C06 were consistent across FSHD patient lines, we tested it in two FSHD1 primary myogenic cell lines, one FSHD1 immortalized line, and one FSHD2 primary myogenic cell line. C06 consistently reduced *DUX4-fl* expression across all tested lines (Figure 4), demonstrating robust activity in multiple cellular contexts.

**Figure 3.**
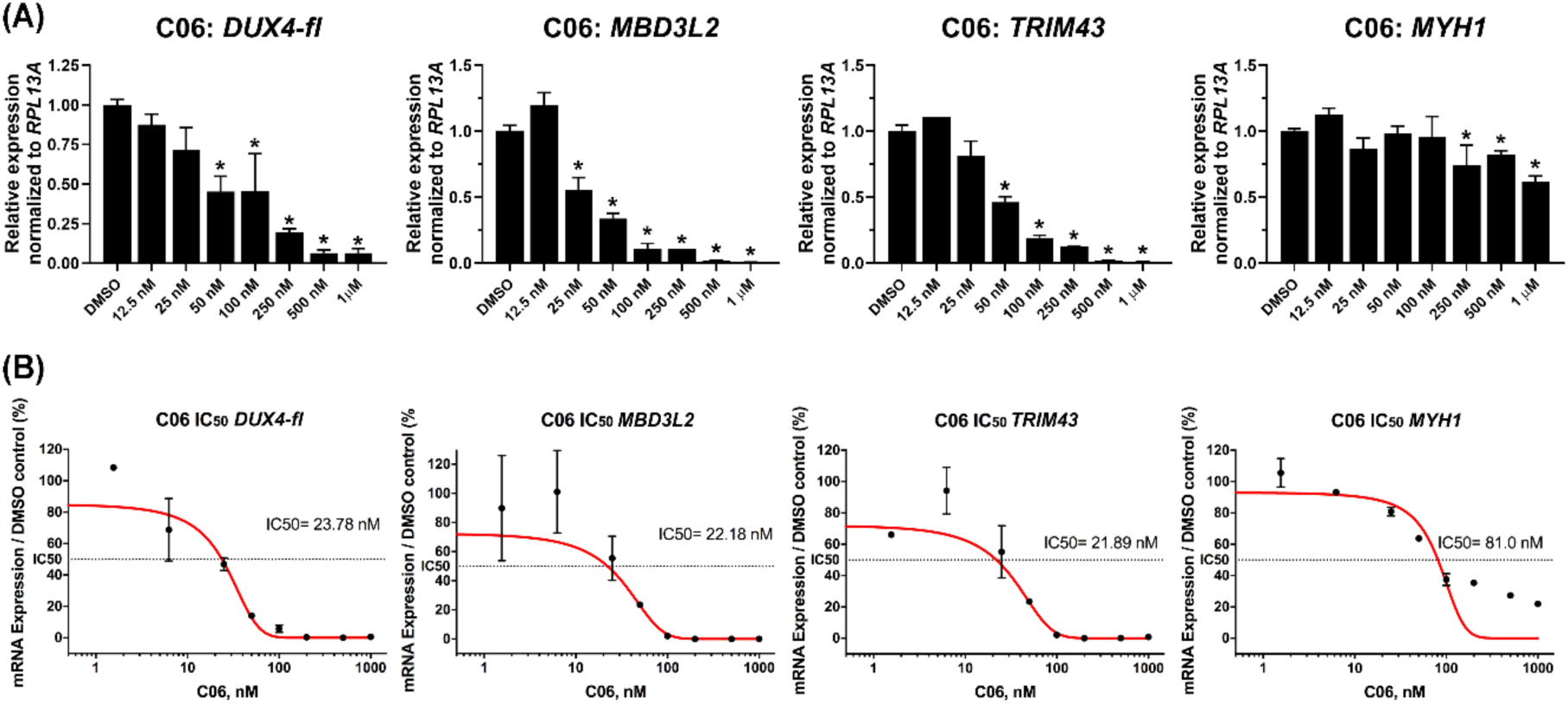
C06 treatment yields specific and dose-dependent suppression of *DUX4-fl* and DUX4-FL targets in FSHD1 myocytes. (**A**) FSHD1 myoblasts (15Ai) were differentiated and treated with C06 at serial concentrations from 12.5 nM to 1 μM. Expression levels of *DUX4-fl,* DUX4-FL target genes *MBD3L2* and *TRIM43,* and the differentiation marker *MYH1* were assessed by RT-qPCR and normalized to *RPL13A*. Data are plotted as the mean ± standard error of the mean (SEM) of at least three independent experiments, with average expression of vehicle-treated cells (DMSO) set to 1. \**p* < 0.05 is from comparing to DMSO. **(B)** IC50 values were calculated for the transcripts in (A) using a nonlinear regression model to fit dose-response curves to the log inhibitor concentration vs. normalized expression levels.

**Figure 4.**
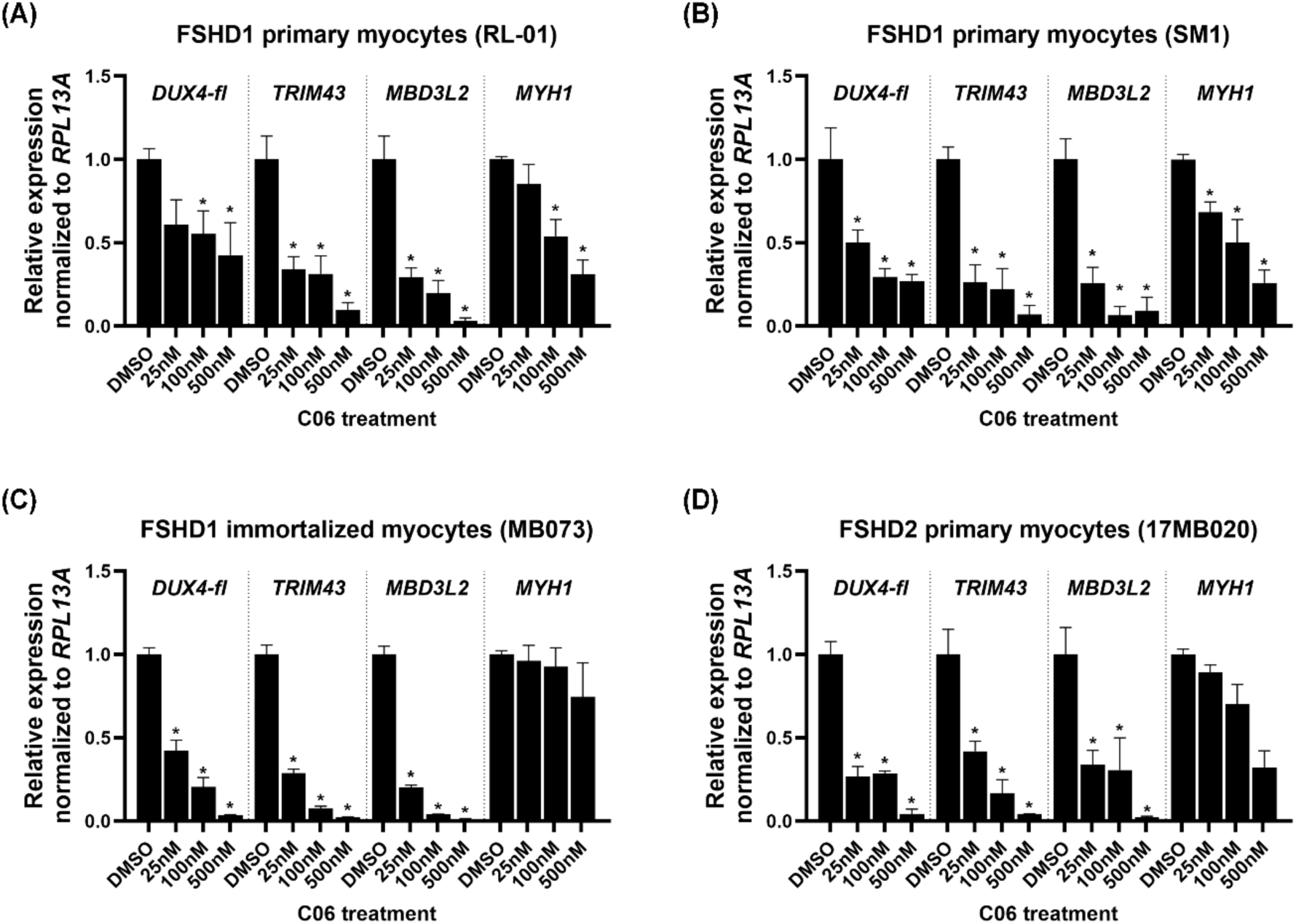
C06 treatment suppresses *DUX4-fl* and DUX4-FL targets in multiple FSHD muscle cell lines. Two primary FSHD1 lines (A-B), one immortalized FSHD1 line (C), and one primary FSHD2 line (D) of patient myoblasts were differentiated and treated with C06 at 25 nM, 100 nM, and 500 nM. Expression levels of *DUX4-fl,* DUX4-FL target genes *MBD3L2* and *TRIM43,* and the differentiation marker *MYH1* were assessed by RT-qPCR and normalized to *RPL13A*. Data are plotted as the mean ± standard error of the mean (SEM) of at least three independent experiments, with average expression of vehicle-treated cells (DMSO) set to 1. \**p* < 0.05 is from comparing to DMSO.

To assess the potential cytotoxicity of C06, we evaluated its effects in primary human dermal fibroblasts. The results indicated that only extremely high concentrations of C06 caused cell death; those below 100 μM did not significantly affect cell viability over time (Figure 5A). Consistent with this, dual viability assays confirmed that C06 treatment did not compromise cell survival (Figure 5B). Collectively, these findings support that C06 is capable of potent and non-toxic *DUX4* suppression, establishing it as our lead compound for further development.

**Figure 5.**
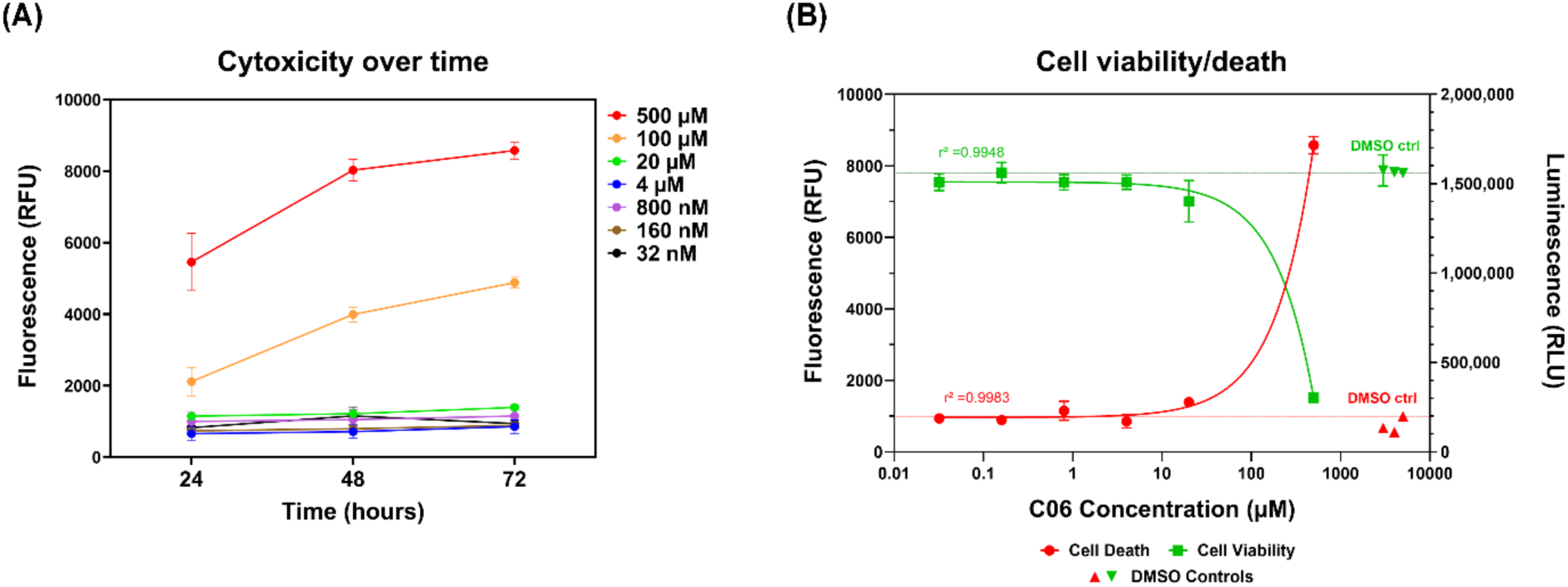
C06 treatment is not cytotoxic. Healthy fibroblasts were treated with C06 at serial concentrations from 32 nM to 500 μM over the indicated timecourse. **(A)** Cytotoxicity was assessed by a fluorometric assay (CellTox™ Green Cytotoxicity Assay). **(B)** Cell survival was assessed by a luminometric assay (CellTiter-Glo® 2.0 Assay). Data are plotted as the mean ± standard error of the mean (SEM) of three independent experiments.

To evaluate the binding of C06 to BAZ1A, molecular docking was performed. Docking of C06 yielded a top scoring pose with a predicted binding score of −5.0 kcal/mol. In the predicted binding mode, C06 forms hydrogen bonding interactions with the N-acetyl lysine binding pocket of BAZ1A, including contacts with residues N1509 and the gatekeeper residue E1515. The hydrogen bond distances were approximately 2 Å with a bent interaction geometry (Figure 6A). The docking results suggested that C06 can be accommodated within the binding pocket of BAZ1A, providing a structural rationale for further experimental validation. To verify the ability of C06 to bind the BAZ1A bromodomain, we performed differential scanning fluorimetry, and found a small, but detectable shift, as indicated by a melting temperature (T_m_) decrease of ∼0.67°C, indicative of binding (Figure 6B). Interestingly, while C06 was predicted to bind the BAZ1A bromodomain pocket, the compound contains elements that can engage kinase ATP pockets as well, namely a pyridine nitrogen that can interact with the hinge motif, a carbonyl that can interact with the aspartate backbone NH from the conserved DFG motif, and a *tert*-butyl group which is positioned to engage the hydrophobic back pocket. Furthermore, a crystal structure of a compound identical to C06 bound to a non-receptor tyrosine kinase, PYK2, was reported which confirms the interactions with the pyridine, carbonyl, and *tert*-butyl moieties (Supplementary Figure S3)^44^. The *tert*-butyl group is critical for kinase binding^32^ since it engages the back pocket (Supplementary Figure S4) while the *tert*-butyl is potentially less critical for BAZ1A binding since docking studies show that it is solvent exposed (see Figure 6A). Therefore, in an attempt to reduce kinase binding while maintaining BAZ1A binding, we synthesized an analog of C06 lacking the *tert*-butyl group (C06-Δt) (Figure 6C). However, differential scanning fluorimetry showed that C06-Δt produced essentially no thermal shift compared to the vehicle control, indicating that the *tert*-butyl group is required for binding of C06 to the BAZ1A bromodomain (Figure 6D). As expected, C06-Δt was also unable to repress *DUX4* and its downstream targets in FSHD myocytes (Figure 6E). Thus, the *tert*-butyl group is required for the ability of C06 to bind BAZ1A and subsequently repress *DUX4* expression.

**Figure 6.**
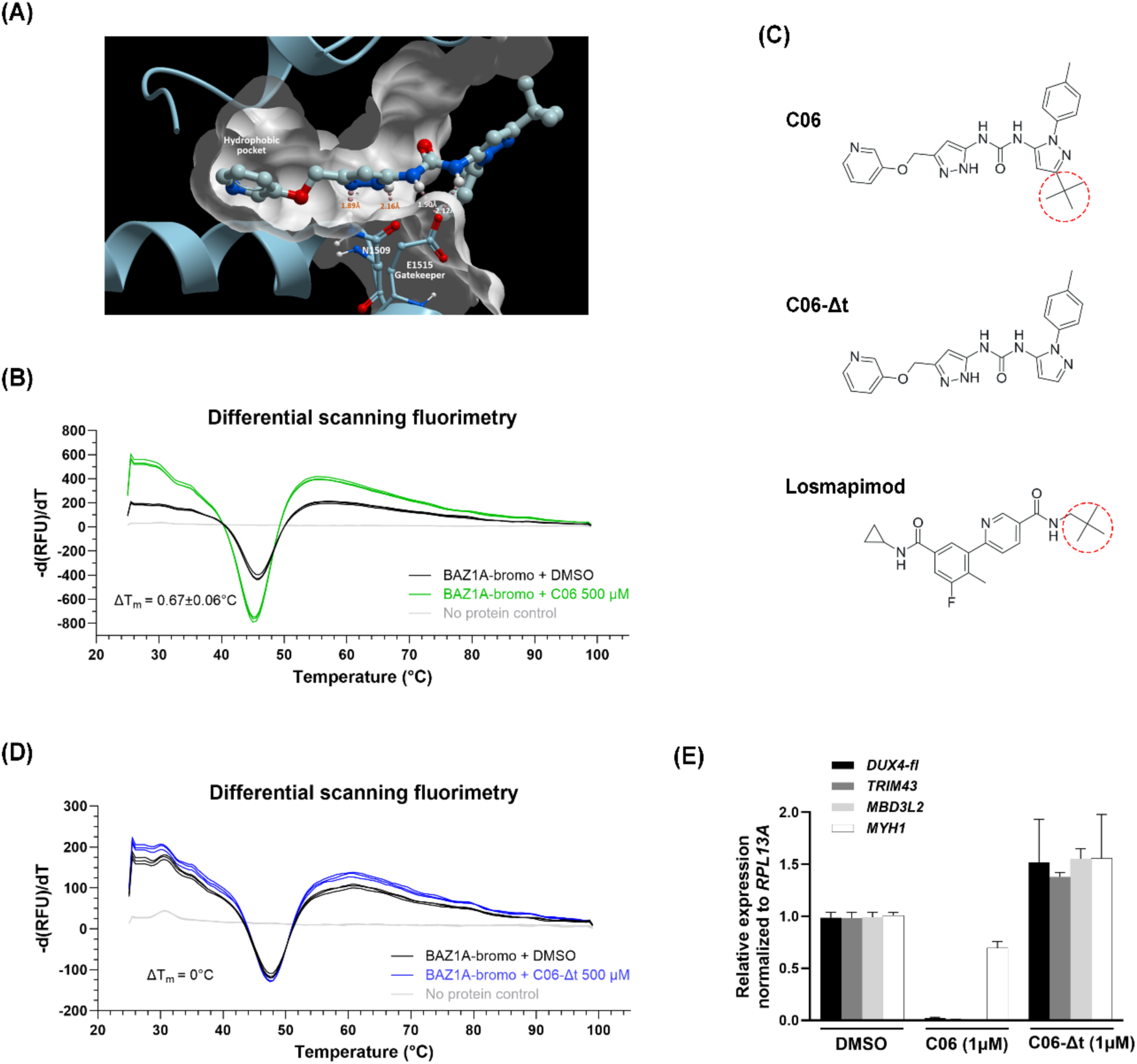
The C06 *tert*-butyl group is required for binding to BAZ1A. (**A**) Predicted structure of C06 bound to the BAZ1A bromodomain. Predicted interactions with residues N1509 and the gatekeeper residue E1515 are shown. The structure was rendered using ICM-Pro ver 3.9-5 (MolSoft LLC). **(B, D)** The thermal stability of the recombinant BAZ1A bromodomain (BAZ1A-bromo) in the presence of 500 µM C06 or C06-Δt was assessed by differential scanning fluorimetry. ΔT_m_ represents the change in melting temperature (T_m_) of BAZ1A-bromo upon compound treatment relative to DMSO. **(C)** Chemical structures of compounds C06, C06-Δt, and the p38α inhibitor losmapimod rendered using Marvin 25.3.3 (ChemAxon Ltd). Red dashed circles indicate the *tert*-butyl group. **(E)** FSHD1 myoblasts (MB073) were differentiated and treated with C06 or C06-Δt at 1 µM. Expression levels of *DUX4-fl,* DUX4-FL target genes *MBD3L2* and *TRIM43,* and the differentiation marker *MYH1* were assessed by RT-qPCR and normalized to *RPL13A*. Data are plotted as the mean ± standard error of the mean (SEM) of three independent experiments, with average expression of vehicle-treated cells (DMSO) set to 1.

To assess the ability of C06 to associate with kinases, we performed a kinome profiling of C06 across a panel of 330 kinases (Supplementary Table S2). Eleven kinases, including p38α, exhibited ≥ 70% inhibition by 0.5 µM C06 (Figure 7A). Many of these kinases are expressed in skeletal muscle and play a role in muscle development, differentiation, or regeneration^24,33–36^. Interestingly, although PYK2 was reported to be inhibited by C06^37^, this kinase only exhibited 37% inhibition by C06 in this assay (Supplementary Table S2).

**Figure 7.**
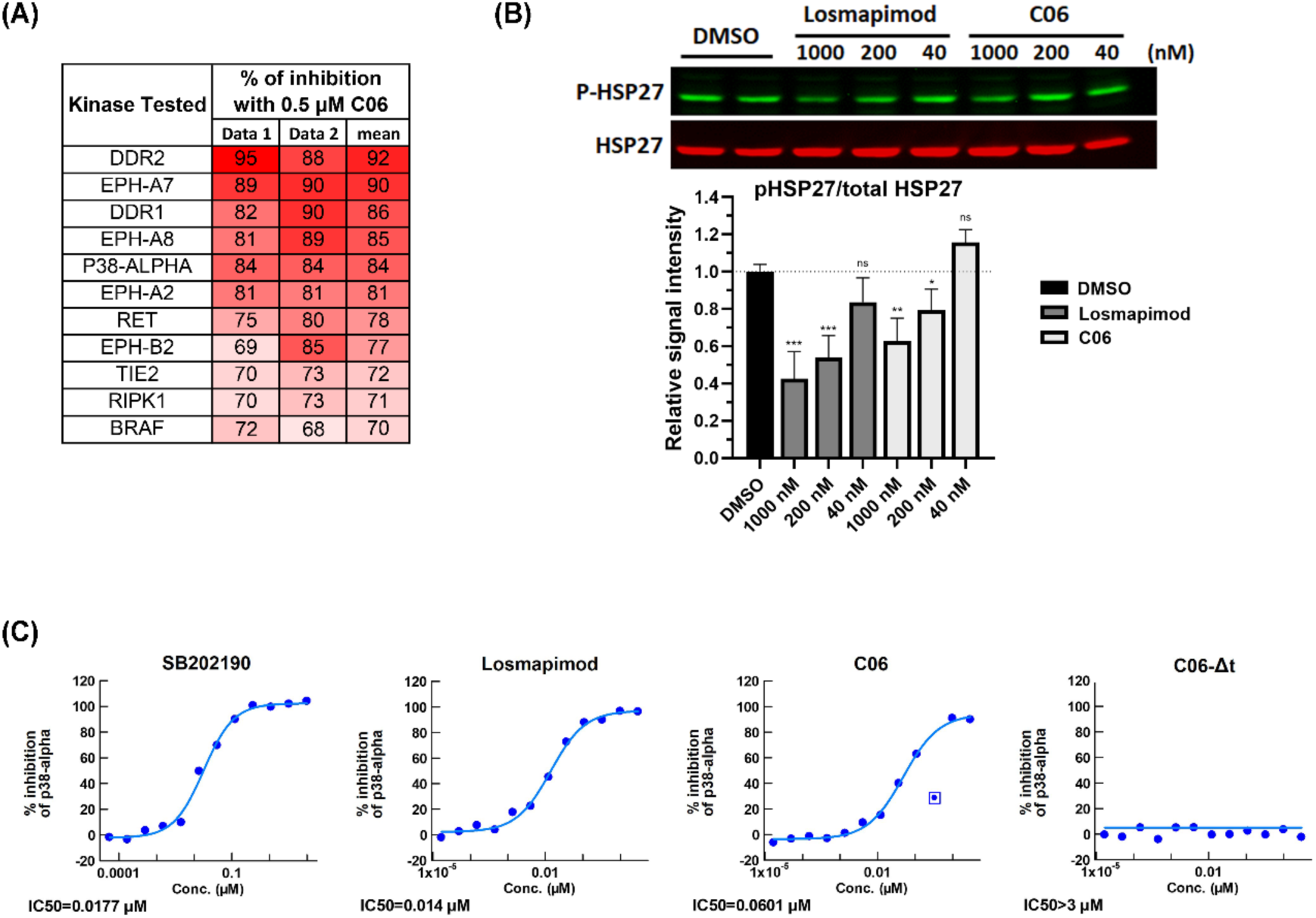
C06 inhibits kinases. **(A)** C06 kinase profile. Listed are kinases exhibiting at least 70% inhibition of activity following treatment with 0.5 µM C06, compared with DMSO control. Values represent the mean of two independent experiments. **(B)** Representative Western blot of phosphorylated heat shock protein 27 (pHSP27) and total HSP27 in FSHD1 myotubes (MB073) treated for 12 hours with losmapimod or C06 at serial concentrations (40-1000 nM). The lower panel shows quantitation of relative intensities. Data are plotted as the mean ± standard error of the mean (SEM) of three independent experiments, with average expression of vehicle-treated cells (DMSO) set to 1. \**p* < 0.05, \*\**p* < 0.01, \*\*\**p* < 0.001 are from comparing to DMSO. The original blots are presented in Supplementary Figure S5. **(C)** Inhibition curves of p38α activity determined by a kinase assay. Losmapimod, C06, and C06-Δt were tested against p38α in a 12-point concentration-response format with 3-fold serial dilutions (0.0169 nM to 3 µM). SB202190 is a positive control for p38 inhibition.

p38α is a serine/threonine mitogen-activated protein kinase (MAPK) that serves as a key regulator of muscle biology^24,25^ and an upstream regulator of *DUX4*^22,38,39^. To determine whether C06 affects the p38α pathway in FSHD myocytes, we measured phosphorylation of heat shock protein 27 (HSP27), a downstream substrate of p38α. We found that C06 inhibits p38α signaling in differentiated FSHD myocytes with an IC50 ∼4.3X higher than losmapimod (Figure 7B-C). By contrast, C06-Δt had no effect on the activity of p38α (Figure 6C). Thus, the *tert*-butyl group of C06 is required for its activity mediated through both BAZ1A and p38α.

## Discussion

Unlike most muscular dystrophies, which are caused by the loss of a factor that plays a critical role in muscle, FSHD is caused by the misexpression of a protein that is toxic in skeletal muscle, due to epigenetic dysregulation of the disease locus. The identification of factors and mechanisms regulating D4Z4 chromatin^16^ has greatly expanded the spectrum of potential therapeutic targets in FSHD. Here we investigated the therapeutic potential of targeting BAZ1A, previously identified as an upstream activator of *DUX4*^26^. We employed an AI-based screening pipeline to identify small molecules predicted to bind the BAZ1A bromodomain, which were subsequently validated using FSHD-specific assays. This led to the identification of C06, a potent suppressor of *DUX4* expression in FSHD myocytes.

Taken together, our results suggest that C06 might function to suppress *DUX4* expression through multiple pathways. The compound was predicted to bind the BAZ1A bromodomain and did exhibit binding in vitro; in a separate study, we have shown that C06 reduces occupancy of BAZ1A at *DUX4* and increases chromatin repression across the *DUX4* locus (Chang et al., manuscript in preparation). However, C06 can also inhibit multiple kinases, including p38α, an established activator of *DUX4.* As a p38α inhibitor, C06 is ∼23% as potent as losmapimod. Interestingly, while even low concentrations of losmapimod led to significant reductions in *MYH1* expression across patient cell lines, C06 had little to no effect on *MYH1* expression at a range of concentrations. Indeed, at low concentrations which still effectively repress *DUX4-fl,* C06 has strikingly few effects on the muscle transcriptome outside of DUX4-FL targets (Chang et al., manuscript in preparation). This strongly suggests that at these lower concentrations, kinase pathways are not disrupted. Considering that most of the kinases exhibiting ≥ 70% inhibition by C06 have a known role in skeletal muscle, this suggests that dose is key to achieving specificity of action. Additionally, C06 may be inhibiting kinases which mediate the pathological effects of DUX4-FL expression in muscle cells, thus allowing a return to a healthier state. For example, in addition to p38α, both the receptor tyrosine kinase RET and the serine/threonine kinase RIPK1 exhibited ≥ 70% inhibition by C06 and have a known association with DUX4. DUX4 upregulates the expression of RET, which is a key regulator of muscle satellite cell proliferation; RET inhibition rescued both the proliferation and differentiation defects in FSHD myoblasts and improved their regenerative capacity^35^. RIPK1 is a key regulator of muscle cell death and inflammation^40^ that mediates DUX4-induced necroptosis^41^. Thus, C06 is likely acting both upstream and downstream of *DUX4* in FSHD muscle cells, which may provide an explanation for its potency. It is possible that modest inhibition of multiple pathways converging on *DUX4* can lead to equal efficacy and greater specificity of action than robust inhibition of a single pathway, as the comparisons between C06 and losmapimod suggest. Studies testing this hypothesis are currently underway.

Our results highlight both the utility and limitations of using AI structural predictions for initial compound screening combined with a function-based (rather than a target-binding) assay. The AI pipeline allowed us to validate a relatively small number of compounds using laborious, but directly relevant assays (*DUX4-fl* and target gene expression) in the relevant cell type (FSHD myocytes). However, while this strategy resulted in the identification of potent *DUX4* suppressors, it did not necessarily identify compounds that bind the intended target with the highest affinity. Thus, our study highlights a critical caveat of AI-based screening approaches, and the need for subsequent target validation. Ultimately, our approach still proved useful, since we were interested in finding a compound that acted as a potent *DUX4* repressor, with the potential for further optimization of specificity and metabolic properties. Preliminary ADME (absorption, distribution, metabolism, elimination) analysis (Supplementary Figure S6 and Supplementary Table S3) revealed that C06 exhibited poor drug-like properties for use in vivo, being rapidly metabolized by liver microsomes. Interestingly, two other hit compounds in our initial screen, A01 and D08, which exhibit much better metabolic properties, demonstrated consistent repression of *DUX4* and its targets, but this repression was not dose-dependent.

Therapeutic development for FSHD remains challenging, and innovative strategies are needed to complement traditional approaches^16^. Here, we demonstrated that an AI-driven, candidate-based screening strategy can efficiently identify small molecules that repress *DUX4* expression in FSHD myocytes. While C06 requires further chemical derivatization to improve its pharmacokinetic profile, the compound represents a useful in vitro tool for potent and specific *DUX4* suppression, and for dissecting the epigenetic regulation of the FSHD locus. What we have learned so far underscores both the promise and current limitations of AI-assisted drug discovery, and emphasizes the need for careful experimental validation of predicted compounds.

## Methods

### Human subjects

The study was conducted according to the guidelines of the Declaration of Helsinki. Muscle biopsy for myoblast isolation was approved by the University of California, Davis Institutional Review Board (IRB #1518564, approved on 18 November 2020). Skin biopsy and primary fibroblast isolation was approved by the University of Nevada, Reno IRB (#1316095, approved on 5 December 2019). Residual surgical tissue from scapular fixation tissue was declared IRB exempt by the University of Nevada, Reno IRB (#1153216, approved 26 February 2018). All qualified subjects provided written informed consent.

### Cell lines and cell cultures

Myogenic cells derived and immortalized from biceps muscle of a patient with FSHD1 (15Ai) were obtained from the Sen Paul D. Wellstone Cooperative Research Center FSHD cell repository^42,43^ and grown as described^44^. Another immortalized FSHD1 myoblast line (MB073) (Fields Center for FSHD and Neuromuscular Research) was obtained from Dr. Stephen Tapscott at the Fred Hutchinson Cancer Center and grown as described^20,45^. Two FSHD1 primary myoblast lines (SM1 and RL-01), and one human dermal fibroblast line (HDF-10064) were established and expanded in the Jones lab at the University of Nevada, Reno School of Medicine, and grown as described^46^. RL-01 was obtained from a needle biopsy of the vastus lateralis from an adult FSHD1 patient (9 D4Z4 repeat units). SM1 was obtained from residual scapular fixation tissue obtained from an early onset FSHD1 patient. HDF-10064 was derived from skin punch biopsy samples from a healthy, non-FSHD subject. Primary FSHD2 myoblasts (17MB020) was obtained from Dr. Rabi Tawil (Fields Center for FSHD and Neuromuscular Research), and grown as described^47^.

For differentiation of 15Ai cells, myoblasts were seeded onto 0.1% gelatin-coated 12-well plates at 1.7 x 10^5^ cells per well and allowed to adhere for 24-72 hours in RoosterNourish™ MSC medium (RoosterBio). Once cultures reached 70-80% confluence, differentiation was induced by replacing the medium with NbActiv4 (Brainbits/TransnetYX Tissue, #Nb4-500) following a wash with 1X phosphate-buffered saline (PBS, Corning^TM^ Cellgro^TM^, #20-031-CV) to remove residual growth factors. Small-molecule compounds were dissolved in dimethyl sulfoxide (DMSO, Sigma-Aldrich, #D1435) and added at the indicated final concentrations at the onset of differentiation. Cells treated with an equivalent volume of DMSO served as vehicle controls. Differentiating myocytes were fed daily with fresh NbActiv4 medium with compounds re-administered at each feeding. To enhance cell viability, horse serum (Sigma-Aldrich, #H1270) was supplemented at a final concentration of 1% after the first 24 hours of differentiation. After four days of differentiation and treatment, cells were harvested for gene expression analysis.

For MB073 immortalized myoblasts and all other primary myoblasts, cells were maintained in Ham’s F10 (Corning™ Cellgro™, #10-070-CV) medium supplemented with 15% fetal bovine serum (FBS) (HyClone), 10 ng/mL rhFGF-basic (PeproTech, 100-18B), and 1 µM dexamethasone (for MB073, Sigma, D4902), or 20% FBS and 1% chick embryo extract (for primary cells) under growth conditions. Differentiation was induced in Dulbecco’s modified Eagle medium (DMEM) (Corning™ Cellgro™, #10-013-CV) containing 2% horse serum (for primary cells) or 1.5% horse serum plus 10 µg/ml insulin and 1.2 mM calcium chloride (for MB073).

Primary human dermal fibroblasts (HDF-10064) were cultured in DMEM medium supplemented with 10% FBS, 1X Non-essential amino acid (Cytivia Hyclone, SH30238) and 1% antibiotics/antimycotics (Corning™ Cellgro™, #30-004-CI). Cells were maintained under standard culture conditions and used for cytotoxicity assays.

### AtomNet-based small-molecule virtual screen

A virtual screen for small-molecule inhibitors of BAZ1A was conducted using Atomwise Inc.’s proprietary AI-based AtomNet® structure-based drug discovery platform^30,48^. The AtomNet model was employed to predict the likelihood of small molecules binding to the bromodomain of BAZ1A (aa 1425-1538). Following prediction, screening results were filtered for drug-like properties. Docking and scoring were performed using the AtomNet model using a library of commercially available compound available from Mcule, and compounds were ranked according to their scores. 72 top-scoring compounds were selected for experimental validation (along with 2 DMSO-only controls). Each compound and control was assigned an anonymized identifier to enable blinded functional screening. Compounds were supplied as 10 mM stock solutions in DMSO, shipped to the laboratory under blinded conditions, and stored at −20 °C. Prior to use, stock solutions were diluted to the desired working concentrations for single-use aliquots.

### RNA extraction, reverse transcription, and quantitative real-time polymerase chain reaction (RT-qPCR)

Total RNAs were extracted from myoblasts or differentiated myotubes using TRIzol Reagent (Ambion Life Technologies, #15596018), followed by phenol-chloroform extraction and purification with RNeasy MinElute Cleanup Kit (QIAGEN, #74204), according to the manufacturer’s instructions. 1-2 μg of total RNAs were reverse transcribed using SuperScript^TM^ III Reverse Transcriptase (Invitrogen, #56575) to generate cDNAs. Quantitative PCR (qPCR) was performed using the Bio-Rad CFX96™ Real-Time System (C1000 Touch™ Thermal Cycler) with SYBR Green detection as previously described^43^. For *DUX4-fl* expression analysis, qPCR reactions were performed with 200 ng of cDNA. For the DUX4-FL target genes *MBD3L2* and *TRIM43*, the myogenic marker *MYH1*, and the reference gene *RPL13A*, qPCR reactions were performed with 20 ng, 20 ng, 5 ng, and 10 ng of cDNA, respectively. Primer sequences used for each gene are listed in Supplementary Table S1.

### Cytotoxicity and cell viability assay

*In vitro* cytotoxicity and cell viability were assessed using the CellTox™ Green Cytotoxicity Assay (Promega, #G8742) and the CellTiter-Glo® 2.0 Assay (Promega, #G9241), following the manufacturer’s instructions. Primary human dermal fibroblasts were plated in a 96-well opaque assay plate with 50 μl of growth media per well and allowed to adhere overnight. Cells were then treated with compound C06 at the indicated concentration in a final volume of 100 μl per well and incubated for 72 hours. For cytotoxicity measurements, CellTox™ Green reagents were added to the wells, mixed by orbital shaking (700–900 rpm) for 1 minute to ensure homogeneity, and incubated for 15 minutes at room temperature in the dark. Fluorescence was measured using a fluorescence plate reader (excitation: 485 nm, emission: 520 nm). Following fluorescence detection, plates were equilibrated to room temperature before the addition of CellTiter-Glo reagents to quantify viable cells based on ATP levels. Luminescence was measured after a 5–10 minute incubation using a luminometer to determine cell viability.

### Protein expression and purification

A DNA fragment encoding the BAZ1A bromodomain (amino acids 1425-1538)^49^ was amplified by PCR from a full-length BAZ1A cDNA template and subcloned into the pET23b vector containing an N-terminal hexa-histidine (6×His) tag. The recombinant construct was transformed into competent E. coli BL21 (DE3) cells (New England Biolabs, #C2527). Transformants were inoculated into 2×YT broth and grown overnight at 37°C. The overnight culture was diluted 1:50 into fresh 2×YT medium and grown for 4 hours until it reached mid-log phase (OD₆₀₀ ≈ 0.6–0.8).

Expression of recombinant protein was induced by addition of 1 mM isopropyl β-D-1-thiogalactopyranoside (IPTG) and cultured for an additional 4 hours at 30°C. Cells were harvested by centrifugation and resuspended in Binding Buffer (20 mM Tris–HCl, pH 8.0, 500 mM NaCl, 5 mM imidazole) supplemented with protease inhibitors (Sigma, #P8340). The suspension was lysed by sonication for 4 minutes on ice using ultrasonic pulses (2 s on, 2 s off, 50% amplitude; Branson Sonifier SFX550). The lysate was clarified by centrifugation at 21,300 × g for 15 minutes, and the supernatant was loaded onto a Talon affinity resin column (Takara, #635502). The column was washed with Wash Buffer (20 mM Tris–HCl, pH 8.0, 500 mM NaCl, 15 mM imidazole), and the His-tagged BAZ1A protein was eluted with Elution Buffer (20 mM Tris–HCl, pH 8.0, 500 mM NaCl, 150 mM imidazole). The purified recombinant proteins were confirmed by 12.5 % SDS–PAGE and visualized using Coomassie Brilliant Blue staining. The purified BAZ1A bromodomain protein was used for *in vitro* thermal shift binding assays.

### Molecular docking

Molecular docking studies were performed using ICM-Pro, version 3.9-5 (MolSoft LLC, San Diego, CA). The crystal structure of apo BAZ1A for the docking studies was obtained from the Protein Data Bank (PDB ID: 5UIY, resolution: 1.69Å). Prior to docking, all crystallographic water molecules were removed. The binding pocket and docking grid were identified and defined using the Pocket Finder tool in ICM-Pro. Energy minimization was performed using the default ICM force field parameters to relieve local steric clashes. Ligand structures were prepared in ICM-Pro, including geometry optimization and assignment of protonation states. Docking poses were ranked based on the ICM docking score.

### Differential scanning fluorimetry (protein thermal shift assay)

Protein thermal shift assays were carried out using GloMelt™ 2.0 Thermal Shift Protein Stability Kit (Biotium, #33027) according to the manufacturer’s protocol. Recombinant BAZ1A bromodomain protein was used at 15-25 μM, and test compounds (C06 and C06-Δt) were added at 500 μM. Fluorescence signals were recorded from 25°C to 95°C at a ramp rate of 0.05°C/sec using a Bio-Red CFX96^TM^ Real-Time System (C1000 Touch™ Thermal Cycler). Raw relative fluorescence units (RFU) against temperature data and the corresponding −d(RFU)/dT, which represents the rate of fluorescence change with increasing temperature, were exported from Bio-Rad CFX Maestro 1.1 software (Bio-Rad Laboratories, version 4.1). Melting temperatures (T_m_) were determined using DSFworld (https://gestwickilab.shinyapps.io/dsfworld/).

### Kinome profiling and p38**α** kinase assay

Kinome profiling was conducted by Nanosyn, Inc. (Santa Clara, CA, USA) using a 330-kinase panel. Compound C06 was tested at a final concentration of 0.5 µM. The complete kinome profiling dataset is provided in Supplementary Table S2. The kinase activity assay for p38α was performed by Nanosyn, Inc. (Santa Clara, CA, USA) using the ADP-Glo platform. The assay was performed with 5 nM p38α enzyme, 130 µM ATP, and 8 µM substrate, followed by incubation for 3 hours. Test compounds were prepared in DMSO and serially diluted in 3-fold steps, yielding final assay concentrations ranging from 3 µM to 0.0169 nM. Each concentration was tested in a single well, with the final DMSO concentration maintained at 1%. The reference inhibitor SB202190 was included as a positive control for p38α inhibition.

### Chemical synthesis of C06 derivative

Compound C06-Δt (lacking the *tert*-butyl group) was designed as an analog of compound C06. The compound was custom-synthesized by WuXi AppTec (Shanghai, China), and its structure and purity were confirmed by proton NMR and LC-MS analysis.

### Western blot

Protein lysates were prepared using radio-immunoprecipitation assay lysis buffer composed of 1% NP-40, 0.5% sodium deoxycholate, 0.1% sodium dodecyl sulfate (SDS), 150 mM NaCl, and 50 mM Tris-Cl (pH 8.0), supplemented with protease/phosphatase inhibitor cocktails (Thermo Fisher Scientific, #78440). Equal amounts of total protein (50 μg) were resolved on 10% SDS-polyacrylamide gel using Tris–glycine–SDS (TGS) buffer (25 mM Tris, 200 mM glycine, 0.1% SDS) and subsequently transferred onto polyvinylidene fluoride membranes using TGS buffer containing 20% methanol. Membranes were blocked for 1 hour at room temperature with 5% (*w/v*) bovine serum albumin (BSA) in Tris-buffered saline with 0.1% Tween-20 (TBST; 50 mM Tris pH 7.5, 150 mM NaCl, 0.1% *v/v* Tween-20), followed by overnight incubation at 4 °C with the following primary antibodies: phospho-HSP27 (Cell Signaling Technology, #2401; 1:1000 dilution), total HSP27 (Cell Signaling Technology, #2402; 1:1000 dilution), and α-tubulin (Sigma-Aldrich, #T6199; 1:10,000 dilution). Bound antibodies were detected using Alexa Fluor™–conjugated secondary antibodies (Invitrogen), and fluorescent signals were visualized with Odyssey Imaging System (LI-COR Biosciences). Signal intensities were quantified using Image Studio Software (version 3.1; LI-COR Biosciences).

### Immunofluorescence staining

Cultured cells were fixed with 2% paraformaldehyde (Electron Microscopy Sciences, #15710) in PBS for 10 min at room temperature, followed by gentle washing with PBS. Fixed cells were permeabilized with 0.25% Triton X-100 diluted in PBS for 10 minutes at room temperature and then blocked with 2% normal donkey serum (Jackson ImmunoResearch, #017-000-121) diluted in PBS for 1 hour at room temperature. Cells were incubated overnight at 4 °C with the primary antibody against myosin heavy chain (Developmental Studies Hybridoma Bank, #MF20) at a concentration of 1–1.5 µg/mL diluted in blocking buffer. Cells were then incubated for 1 hour at room temperature with secondary antibody (Jackson ImmunoResearch, #715-586-150; 1:500 dilution) diluted in PBS, followed by nuclear staining with DAPI (1 µg/mL in PBS) for 15 minutes at room temperature. Fluorescence images were acquired using a Leica DMi8 fluorescence microscope equipped with appropriate filter sets and Leica LAS X software. For each well, 10 images were captured from at least five non-overlapping regions. Quantification of fusion index was performed using ImageJ software by calculating the percentage of nuclei within myosin heavy chain–positive cells relative to the total number of nuclei in each image.

### Statistics

Experiments in cultured cells were performed using at least three biological replicates, and data was analyzed using an unpaired, two-tailed Student’s t-test performed with GraphPad Prism software version 10.4.2 (GraphPad Software, Inc., La Jolla, CA, USA).

## Data availability

The authors declare that the data supporting the findings of this study are available within the main manuscript and its Supplementary Information files.

## Acknowledgements

The authors thank Lei Zhang for technical assistance.

## Author Contributions

NC, CLH, TIJ, and PLJ designed the study, analyzed data, and wrote the manuscript. NC, CLH, and HPM performed experiments. TO facilitated and conducted the AtomNet-based small-molecule virtual screen. WT provided chemical structure and modeling expertise. BB provided muscle biopsy and clinical expertise. All authors edited and approved the final version of the manuscript.

