## Supplementary for "Identification of compounds that repress DUX4 expression in facioscapulohumeral muscular dystrophy"

### Supplementary Figure S1

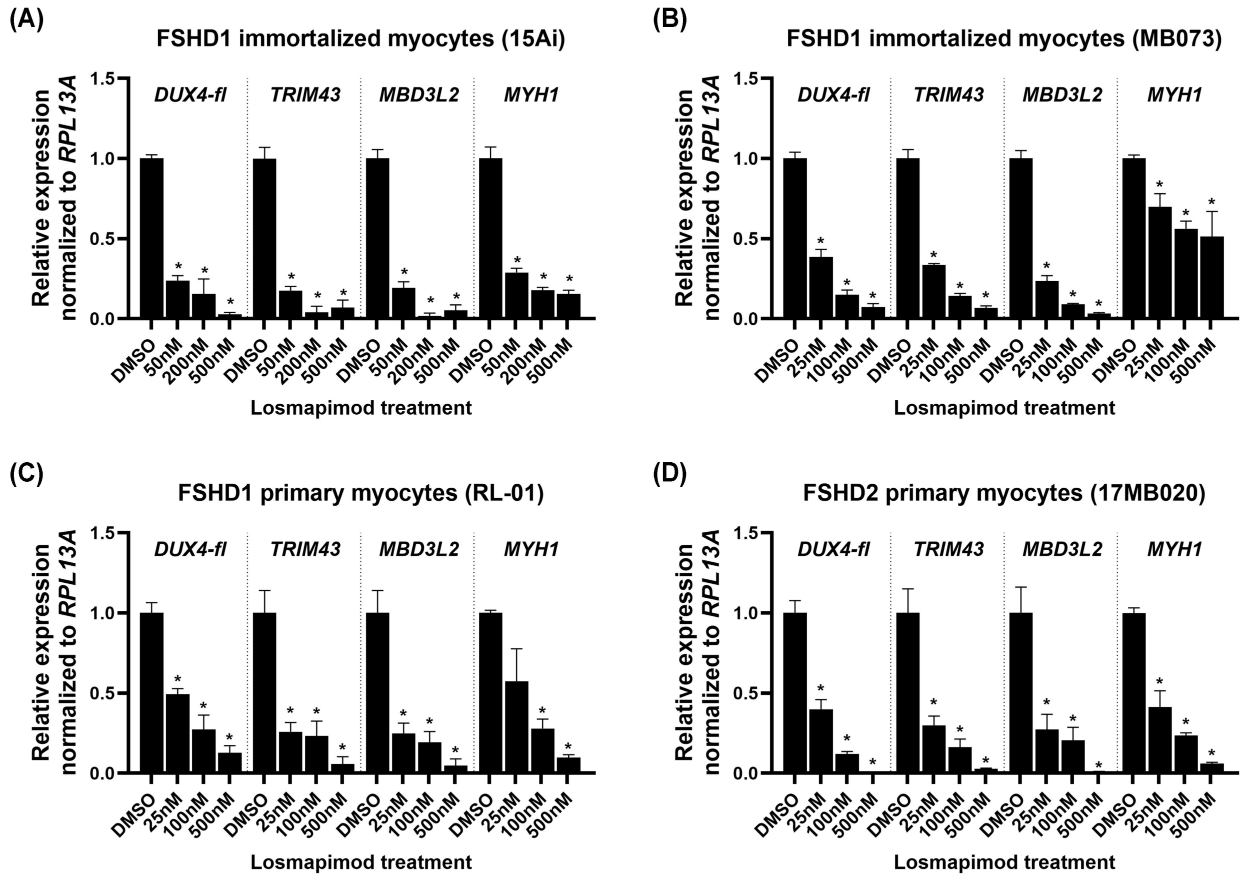


---

**Supplementary Figure S1. Losmapimod treatment suppresses *DUX4-fl*, DUX4-FL targets, and *MYH1* in multiple FSHD muscle cell lines.**

Two immortalized FSHD1 lines (A-B), one primary FSHD1 line (C), and one primary FSHD2 line (D) of patient myoblasts were differentiated and treated with losmapimod at 25 nM (or 50 nM for 15Ai cells), 100 nM, and 500 nM. Expression levels of *DUX4-fl,* DUX4-FL target genes *MBD3L2* and *TRIM43,* and the differentiation marker *MYH1* were assessed by RT-qPCR and normalized to *RPL13A*. Data are plotted as the mean ± standard error of the mean (SEM) of at least three independent experiments, with average expression of vehicle-treated cells (DMSO) set to 1. *$p < 0.05$ is from comparing to DMSO.

### Supplementary Figure S2

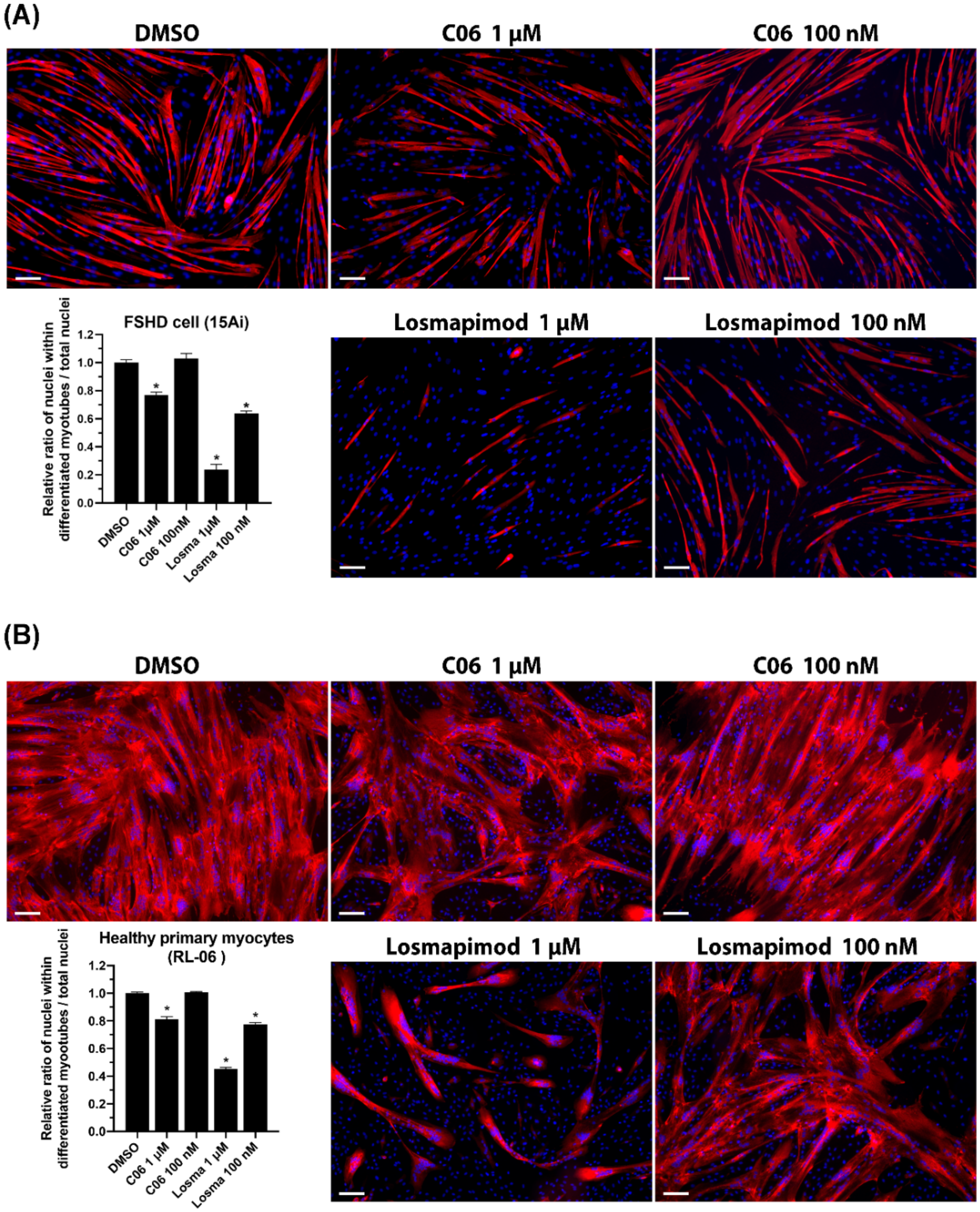


**Supplementary Figure S2. Effects of C06 and losmapimod on myogenic differentiation in FSHD1 and healthy myocytes.**

**(A)** Immunofluorescence staining of differentiated FSHD1 myocytes (15Ai) treated with DMSO, C06 (1 μM or 100 nM), or losmapimod (1 μM or 100 nM) for 4 days. MYH1 was detected by immunostaining (red), and nuclei were counterstained with DAPI (blue). **(B)** Immunofluorescence staining of differentiated healthy primary myocytes (RL-06) under the same treatment conditions. MYH1 (red) and DAPI (blue) staining are shown. Quantification of the ratio of nuclei within MYH1-positive areas is shown (mean ± SEM, *$p < 0.05$ compared with DMSO). Scale bar = 100 μm.

### Supplementary Figure S3.

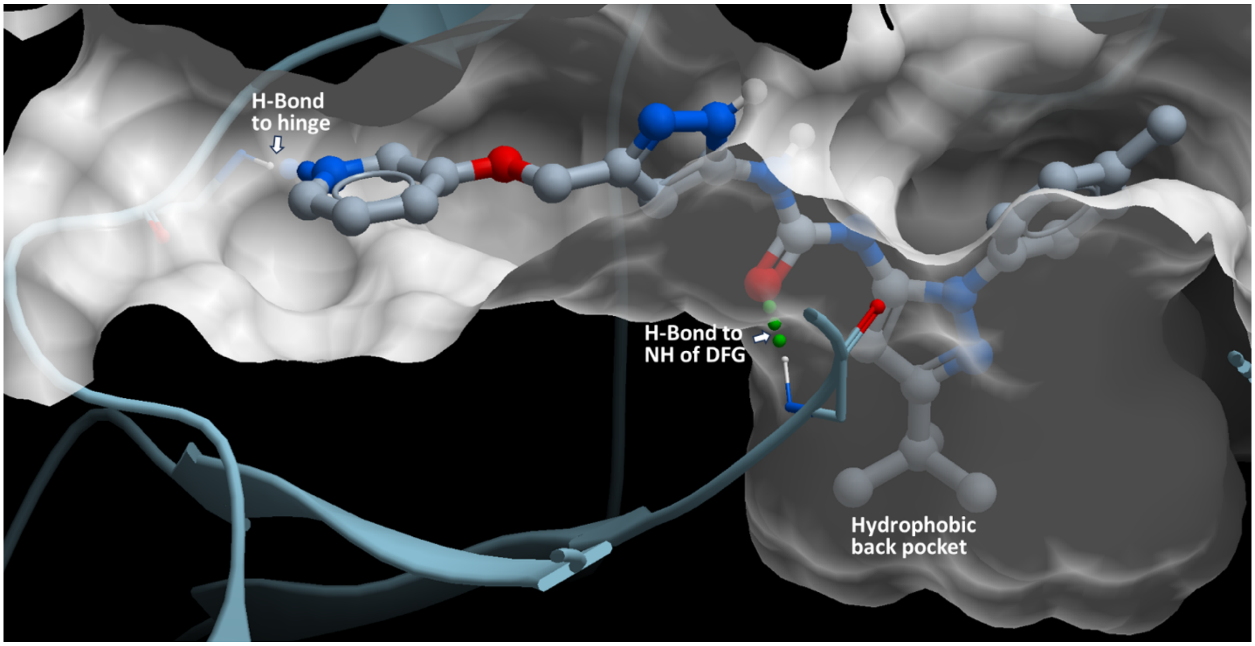


**Supplementary Figure S3. Crystal structure of C06 bound to PYK2.**

Crystal structure of a compound identical to C06 bound to the PYK2 non-receptor tyrosine kinase, reported by Han et al[1]. Interactions with the pyridine, carbonyl, and *tert*-butyl moieties of C06 are shown. The structure was rendered from the PDB file (PDB: 3FZT, resolution 2.00Å) reported by Han et al[1]. using the ICM Browser (Molsoft).

### Supplementary Figure S4

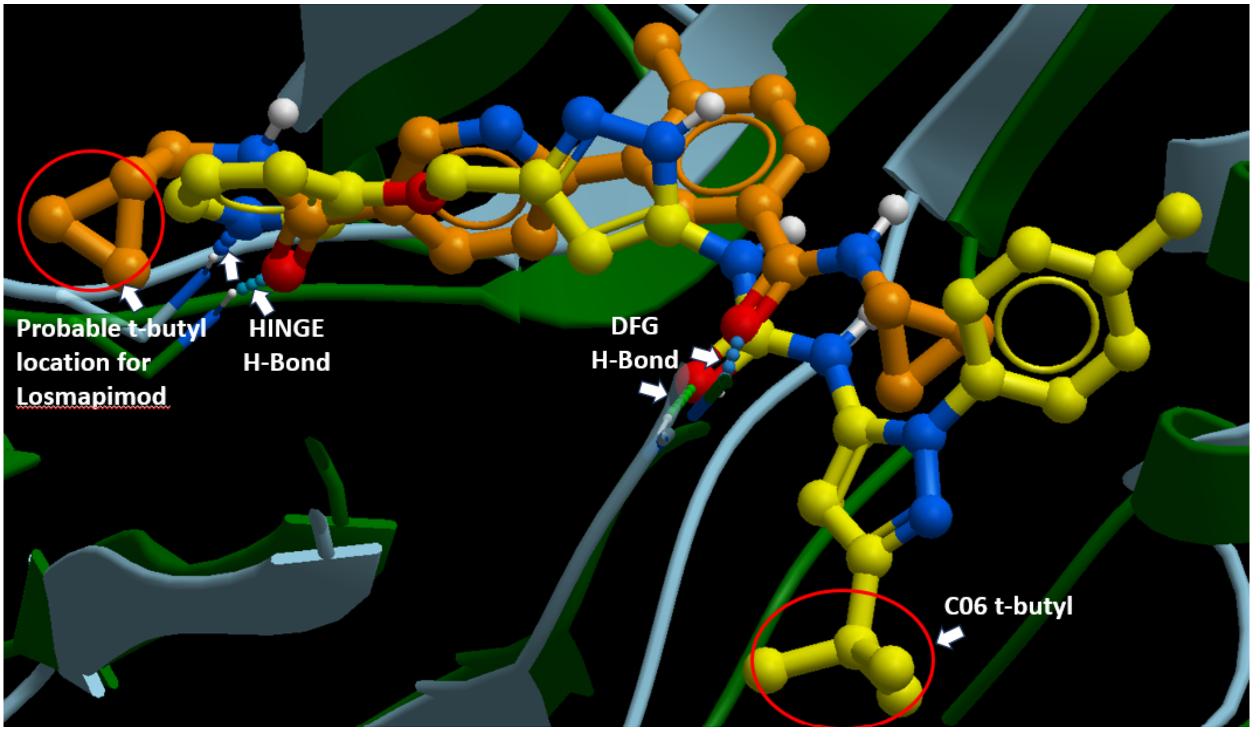


**Supplementary Figure S4. Structural comparison of C06 and losmapimod binding modes.**

Overlay of C06 (yellow; PDB: 3FZT) and a comparator crystal structure representing the binding mode of losmapimod (brown; PDB: 3IPH) in the kinase active site. The *tert*-butyl group of losmapimod is positioned near the mouth of the binding pocket in a solvent-exposed region, whereas the *tert*-butyl group of C06 extends into the deeper back pocket. Key hydrogen-bond (H-Bond) interactions with the hinge region and the DFG motif are indicated. Structural rendering was performed using ICM Browser (Molsoft).

### Supplementary Figure S5

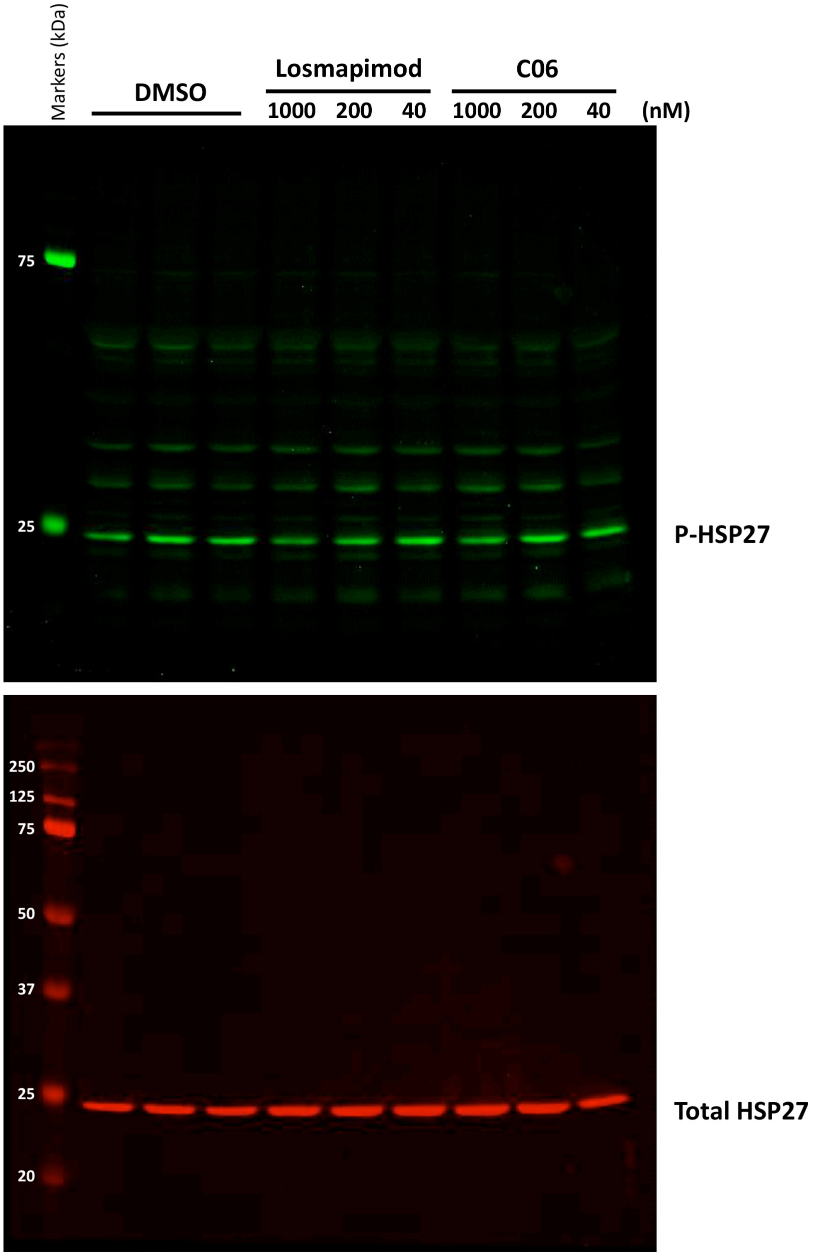


**Supplementary Figure S5. Original western blots corresponding to Figure 7B.**

Uncropped western blot images for phosphorylated HSP27 (P-HSP27) and total HSP27 corresponding to the data shown in Figure 7B. Cells were treated with DMSO or the indicated concentrations of losmapimod or C06 (1000, 200, and 40 nM). P-HSP27 (green, 800 nm channel) and total HSP27 (red, 700 nm channel) were detected by double blotting on the same membrane. Molecular weight marker corresponds to Precision Plus Protein Dual Color Standards (Bio-Rad, #1610374).

### Supplementary Figure S6

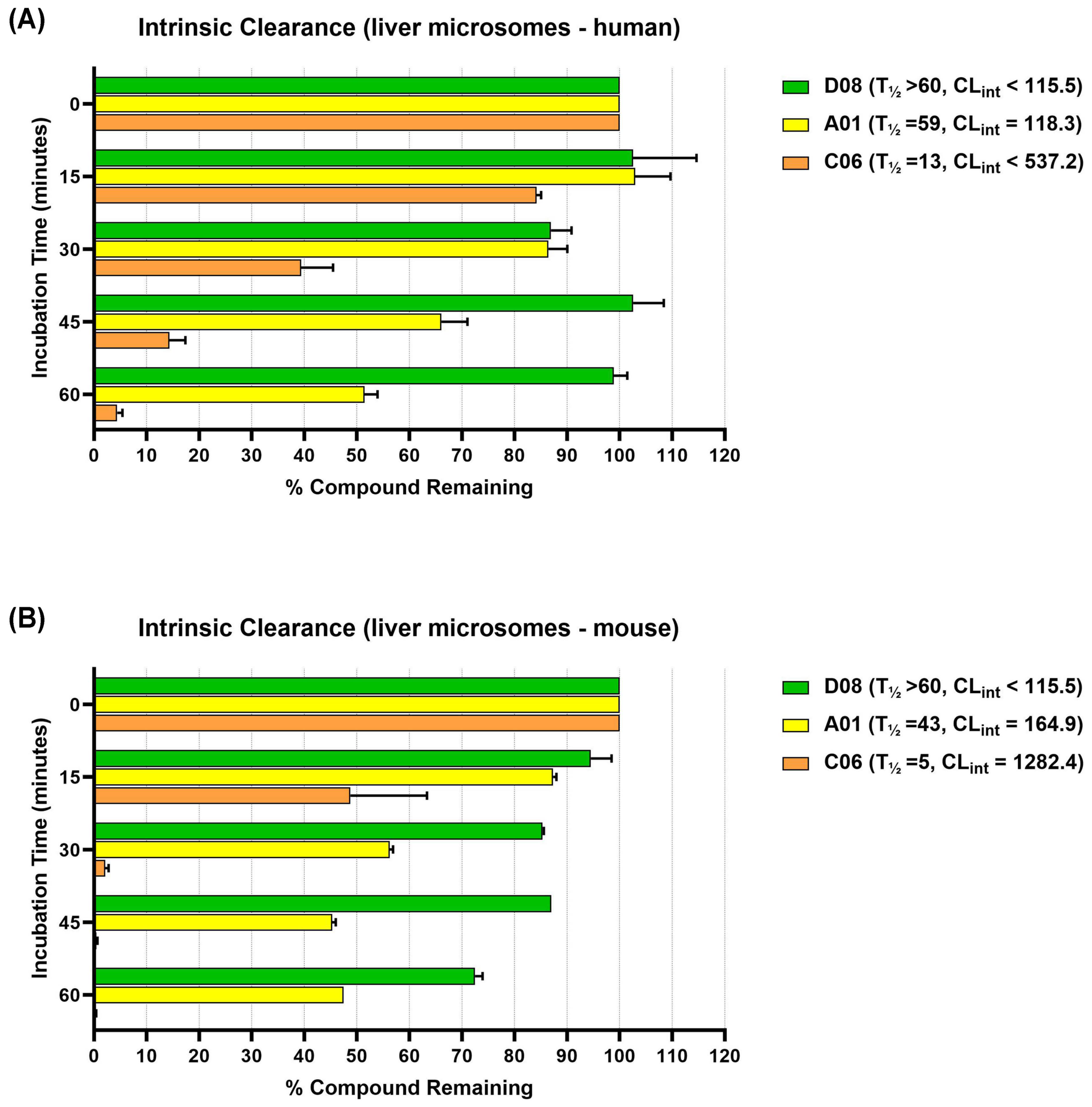


**Supplementary Figure S6. ADME (absorption, distribution, metabolism, elimination) analysis.**

Three compounds from the AI screen that effectively repress *DUX4-fl* and DUX4-FL targets (D08, A01, and C06) were tested for intrinsic clearance by liver microsomes using **(A)** human liver microsomes or **(B)** mouse liver microsomes. Data is shown as percent of the parent compound remaining at 15, 30, 45, and 60 minutes relative to that at the beginning. Data represent the mean of two experiments. The half-life ($T_{1/2}$) was estimated from the slope of the initial linear range of the logarithmic curve of compound remaining (%) vs. time, assuming first-order kinetics. The value of intrinsic clearance ($CL_{int}$, in µL/min/mg for microsomes) was calculated according to the following formula:

$$CL_{int} = \frac{0.693}{T_{1/2} * (\text{mg protein/µL or million cells/µL or pmol CYP isoyme/µL})}$$

**Supplementary Table S1. Oligonucleotide primers**

| Primer name | Sequence |
|---|---|
| *DUX4-fl*-F (prePCR) | 5′-GCTCTGCTGGAGGAGCTTTAGGA-3′ |
| *DUX4-fl*-R (prePCR) | 5′-CGCACTGCTCGCAGGTCTGCWGGT-3′ |
| *DUX4-fl*-nested-F | 5′-AGCTTTAGGACGCGGGGTTGGGAC-3′ |
| *DUX4-fl*-nested-R | 5′-GCAGGTCTGCWGGTACCTGG-3′ |
| *MBD3L2*-F | 5′-GCGTTCACCTCTTTTCCAAG-3′ |
| *MBD3L2*-R | 5′-GCCATGTGGATTTCTCGTTT-3′ |
| *TRIM43*-F | 5′-ACCCATCACTGGACTGGTGT-3′ |
| *TRIM43*-R | 5′-CACATCCTCAAAGAGCCTGA-3′ |
| *RPL13A*-F | 5′-AACCTCCTCCTTTTCCAAGC-3′ |
| *RPL13A*-R | 5′-GCAGTACCTGTTTAGCCACGA-3′ |

### Supplementary Table S2.  Kinome profiling results

| Kinase Tested | Compound Concentration (µM) | Compound Name | % Inhibition | | % Inhibition (mean) | Reference IC50 values | | |
|---|---|---|---|---|---|---|---|---|
| | | | Point 1 | Point 2 | | Staurosporine IC50 (µM) | SB202190 IC50 (µM) | Wortmannin IC50 (µM) |
| DDR2 | 0.5 | C06 | 95 | 88 | 92 | 0.00061 | 1.58110 | >10 |
| EPH-A7 | 0.5 | C06 | 89 | 90 | 90 | 0.19896 | >10 | >10 |
| DDR1 | 0.5 | C06 | 82 | 90 | 86 | 0.00119 | 0.35897 | >10 |
| EPH-A8 | 0.5 | C06 | 81 | 89 | 85 | 0.10001 | >10 | >10 |
| P38-ALPHA | 0.5 | C06 | 84 | 84 | 84 | >10 | 0.00887 | >10 |
| EPH-A2 | 0.5 | C06 | 81 | 81 | 81 | 0.27399 | >10 | >10 |
| RET | 0.5 | C06 | 75 | 80 | 78 | 0.00120 | >10 | >10 |
| EPH-B2 | 0.5 | C06 | 69 | 85 | 77 | 0.03301 | >10 | >10 |
| TIE2 | 0.5 | C06 | 70 | 73 | 72 | 0.18741 | >10 | >10 |
| RIPK1 | 0.5 | C06 | 70 | 73 | 71 | 0.44282 | >10 | >10 |
| BRAF | 0.5 | C06 | 72 | 68 | 70 | 0.00555 | 1.24514 | >10 |
| PDGFR-ALPHA | 0.5 | C06 | 67 | 68 | 67 | <0.000128 | >10 | >10 |
| FLT-3 | 0.5 | C06 | 65 | 62 | 64 | <0.000128 | >10 | >10 |
| TRKC | 0.5 | C06 | 52 | 55 | 53 | 0.00091 | >10 | >10 |
| P38-BETA | 0.5 | C06 | 54 | 51 | 53 | >10 | 0.13076 | >10 |
| ARG | 0.5 | C06 | 45 | 59 | 52 | 0.07442 | >10 | >10 |
| MUSK | 0.5 | C06 | 42 | 44 | 43 | 0.00067 | >10 | >10 |
| EPH-B4 | 0.5 | C06 | 31 | 50 | 41 | 1.01311 | >10 | >10 |
| MNK2 | 0.5 | C06 | 37 | 38 | 38 | 0.01453 | >10 | >10 |
| PYK2 | 0.5 | C06 | 38 | 37 | 37 | 0.01320 | >10 | >10 |
| PDGFR-BETA | 0.5 | C06 | 32 | 42 | 37 | <0.000128 | >10 | >10 |
| CRAF | 0.5 | C06 | 33 | 33 | 33 | >10 | 0.58838 | >10 |
| TRKB | 0.5 | C06 | 35 | 30 | 32 | 0.00094 | >10 | >10 |
| EPH-B1 | 0.5 | C06 | 30 | 33 | 32 | 0.35563 | >10 | >10 |
| YES | 0.5 | C06 | 25 | 32 | 28 | 0.00312 | >10 | >10 |
| EPH-A6 | 0.5 | C06 | 7 | 49 | 28 | 0.03551 | 4.76853 | >10 |
| EPH-B3 | 0.5 | C06 | 22 | 33 | 28 | >10 | >10 | >10 |
| BRK | 0.5 | C06 | 23 | 25 | 24 | 0.82074 | 3.16521 | >10 |
| TYK2 | 0.5 | C06 | 25 | 20 | 23 | <0.000128 | >10 | >10 |
| HCK | 0.5 | C06 | 19 | 25 | 22 | 0.00148 | >10 | >10 |
| FMS | 0.5 | C06 | 19 | 23 | 21 | 0.00022 | >10 | >10 |
| LYNA | 0.5 | C06 | 15 | 27 | 21 | 0.00194 | >10 | >10 |
| ATR | 0.5 | C06 | 17 | 25 | 21 | >10 | >10 | 3.43513 |
| SRC | 0.5 | C06 | 15 | 27 | 21 | 0.00199 | >10 | >10 |
| RSK3 | 0.5 | C06 | 18 | 22 | 20 | <0.000128 | >10 | >10 |
| LOK | 0.5 | C06 | 26 | 14 | 20 | <0.000128 | >10 | >10 |
| FLT-1 | 0.5 | C06 | 16 | 21 | 19 | 0.00800 | >10 | >10 |
| MAP2K7 | 0.5 | C06 | 38 | -2 | 18 | 0.04783 | >10 | >10 |
| FAK | 0.5 | C06 | 11 | 24 | 17 | 0.03764 | >10 | >10 |
| PI3K-GAMMA | 0.5 | C06 | 9 | 25 | 17 | 9.97081 | >10 | 0.00032 |
| FGR | 0.5 | C06 | 29 | 5 | 17 | 0.00351 | >10 | >10 |
| ABL1 | 0.5 | C06 | 0 | 34 | 17 | 0.13721 | >10 | >10 |
| TNIK | 0.5 | C06 | 16 | 15 | 16 | 0.00061 | 7.76309 | >10 |
| LYNB | 0.5 | C06 | 10 | 20 | 15 | 0.00293 | >10 | >10 |
| KDR | 0.5 | C06 | -5 | 35 | 15 | 0.00436 | >10 | >10 |
| RON | 0.5 | C06 | 15 | 14 | 15 | 0.13811 | >10 | >10 |
| KIT | 0.5 | C06 | 16 | 14 | 15 | 0.01189 | >10 | >10 |
| EPH-A5 | 0.5 | C06 | -6 | 34 | 14 | 0.06542 | >10 | >10 |
| FGFR1 | 0.5 | C06 | 11 | 18 | 14 | 0.00793 | >10 | >10 |
| EGFR | 0.5 | C06 | 20 | 8 | 14 | 1.31829 | >10 | >10 |
| ERB-B4 | 0.5 | C06 | 13 | 15 | 14 | 0.74760 | >10 | >10 |
| FLT-4 | 0.5 | C06 | 12 | 16 | 14 | 0.00105 | >10 | >10 |
| FGFR4 | 0.5 | C06 | 16 | 11 | 14 | 0.48975 | >10 | >10 |
| JAK3 | 0.5 | C06 | 7 | 20 | 14 | <0.000128 | >10 | >10 |
| MET | 0.5 | C06 | 17 | 10 | 13 | 0.37001 | >10 | >10 |

| | | | | | | | | |
|---|---|---|---|---|---|---|---|---|
| PI3K-ALPHA | 0.5 | C06 | 9 | 17 | 13 | >10 | >10 | 0.00025 |
| CSK | 0.5 | C06 | 10 | 15 | 12 | 0.20783 | >10 | >10 |
| EPH-A3 | 0.5 | C06 | -16 | 40 | 12 | 0.01759 | >10 | >10 |
| SLK | 0.5 | C06 | 8 | 15 | 12 | <0.000128 | >10 | >10 |
| ITK | 0.5 | C06 | 12 | 11 | 12 | 0.00517 | >10 | >10 |
| SRMS | 0.5 | C06 | 18 | 5 | 12 | 8.19732 | >10 | >10 |
| TNK2 | 0.5 | C06 | 12 | 11 | 11 | 0.00216 | >10 | >10 |
| AKT2 | 0.5 | C06 | 6 | 16 | 11 | 0.02658 | >10 | >10 |
| P38-GAMMA | 0.5 | C06 | 16 | 6 | 11 | 0.59029 | >10 | >10 |
| SYK | 0.5 | C06 | 9 | 13 | 11 | 0.00047 | >10 | >10 |
| FGFR2 | 0.5 | C06 | 12 | 9 | 11 | 0.00268 | >10 | >10 |
| MAP4K5 | 0.5 | C06 | 13 | 8 | 10 | 0.00014 | >10 | >10 |
| MAP4K4 | 0.5 | C06 | 17 | 3 | 10 | <0.000128 | >10 | >10 |
| TXK | 0.5 | C06 | 15 | 5 | 10 | 0.05646 | >10 | >10 |
| AXL | 0.5 | C06 | 3 | 15 | 9 | 0.00149 | >10 | >10 |
| IGF1R | 0.5 | C06 | 5 | 12 | 9 | 0.02636 | >10 | >10 |
| TTK | 0.5 | C06 | 12 | 5 | 8 | 0.13024 | >10 | >10 |
| ROS | 0.5 | C06 | 11 | 6 | 8 | <0.000128 | >10 | >10 |
| TYRO3 | 0.5 | C06 | 12 | 4 | 8 | 0.00108 | >10 | >10 |
| AKT1 | 0.5 | C06 | 11 | 5 | 8 | 0.00971 | >10 | >10 |
| TRKA | 0.5 | C06 | 9 | 6 | 8 | 0.00237 | >10 | >10 |
| TNK1 | 0.5 | C06 | 3 | 11 | 7 | 0.00041 | >10 | >10 |
| MSK2 | 0.5 | C06 | 15 | -1 | 7 | 0.00383 | >10 | >10 |
| LCK | 0.5 | C06 | 2 | 12 | 7 | 0.00247 | >10 | >10 |
| ERB-B2 | 0.5 | C06 | 11 | 2 | 7 | 0.57467 | >10 | >10 |
| SGK2 | 0.5 | C06 | 9 | 4 | 6 | 0.08926 | >10 | >10 |
| TAK1-TAB1 | 0.5 | C06 | -10 | 23 | 6 | 0.03290 | >10 | >10 |
| CLK1 | 0.5 | C06 | 1 | 12 | 6 | 0.00275 | >10 | 1.14316 |
| SPHK1 | 0.5 | C06 | 7 | 5 | 6 | >10 | >10 | >10 |
| MER | 0.5 | C06 | 5 | 7 | 6 | 0.07643 | >10 | >10 |
| FGFR3 | 0.5 | C06 | 7 | 4 | 6 | 0.41618 | >10 | >10 |
| EPH-A1 | 0.5 | C06 | 5 | 6 | 6 | 0.04744 | >10 | >10 |
| TAOK2 | 0.5 | C06 | 12 | 0 | 6 | 0.01079 | >10 | >10 |
| TEC | 0.5 | C06 | 4 | 4 | 4 | 0.08132 | >10 | >10 |
| LTK | 0.5 | C06 | 0 | 8 | 4 | 0.00233 | >10 | >10 |
| FES | 0.5 | C06 | -4 | 11 | 4 | 0.00238 | >10 | >10 |
| IRR | 0.5 | C06 | 2 | 4 | 3 | 0.54753 | >10 | >10 |
| ZAP70 | 0.5 | C06 | 3 | 3 | 3 | 0.00218 | >10 | >10 |
| RSK2 | 0.5 | C06 | 0 | 6 | 3 | <0.000128 | >10 | >10 |
| LZK | 0.5 | C06 | 8 | -2 | 3 | 0.68856 | >10 | >10 |
| MST1 | 0.5 | C06 | -1 | 4 | 2 | <0.000128 | >10 | >10 |
| PKC-BETA1 | 0.5 | C06 | 6 | -3 | 2 | <0.000128 | 1.37659 | 3.27885 |
| ATM | 0.5 | C06 | -5 | 8 | 2 | >10 | >10 | 0.03369 |
| IRAK1 | 0.5 | C06 | 4 | -1 | 1 | 0.09995 | >10 | >10 |
| JAK1 | 0.5 | C06 | 2 | 0 | 1 | 0.00119 | >10 | >10 |
| MAP4K1 | 0.5 | C06 | 4 | -2 | 1 | <0.000128 | >10 | >10 |
| PI3K-DELTA | 0.5 | C06 | -9 | 10 | 0 | >10 | >10 | 0.00072 |
| MRCKA | 0.5 | C06 | 0 | 0 | 0 | 0.00084 | >10 | >10 |
| PKC-ALPHA | 0.5 | C06 | -1 | 1 | 0 | <0.000128 | >10 | 3.31745 |
| NEK4 | 0.5 | C06 | 3 | -5 | -1 | >10 | >10 | >10 |
| P38-DELTA | 0.5 | C06 | -4 | 1 | -1 | 2.02836 | >10 | >10 |
| SPHK2 | 0.5 | C06 | -9 | 7 | -1 | >10 | >10 | >10 |
| CK1-DELTA | 0.5 | C06 | -2 | -1 | -2 | >10 | 0.16992 | >10 |
| PAK4 | 0.5 | C06 | -3 | -1 | -2 | 0.00511 | >10 | >10 |
| PRKD3 | 0.5 | C06 | -5 | 1 | -2 | 0.00091 | >10 | >10 |
| MST2 | 0.5 | C06 | -4 | -1 | -2 | 0.00013 | >10 | >10 |
| DYRK1B | 0.5 | C06 | 2 | -7 | -3 | 0.00471 | >10 | >10 |
| RIPK2 | 0.5 | C06 | -3 | -3 | -3 | 0.44278 | 0.04377 | >10 |
| PDK1 | 0.5 | C06 | -3 | -2 | -3 | 0.00416 | >10 | >10 |
| BMX | 0.5 | C06 | -14 | 8 | -3 | 0.00793 | >10 | >10 |
| NEK3 | 0.5 | C06 | 1 | -7 | -3 | >10 | >10 | 3.75978 |

| | | | | | | | | |
|---|---|---|---|---|---|---|---|---|
| PAK3 | 0.5 | C06 | -3 | -3 | -3 | 0.00126 | >10 | >10 |
| ULK3 | 0.5 | C06 | -7 | 1 | -3 | 0.00066 | >10 | >10 |
| ALK | 0.5 | C06 | -7 | 1 | -3 | 0.00117 | >10 | >10 |
| RSK4 | 0.5 | C06 | -2 | -5 | -3 | <0.000128 | >10 | >10 |
| SGK1 | 0.5 | C06 | -5 | -1 | -3 | 0.02682 | >10 | >10 |
| PIM-1-KINASE | 0.5 | C06 | 0 | -7 | -4 | 0.00246 | >10 | 0.21747 |
| SSTK | 0.5 | C06 | -4 | -3 | -4 | >10 | >10 | >10 |
| CDK5-P25 | 0.5 | C06 | -4 | | -4 | 0.00154 | >10 | >10 |
| FYN | 0.5 | C06 | -9 | 1 | -4 | 0.00645 | >10 | >10 |
| CK2 | 0.5 | C06 | -3 | -5 | -4 | 9.89638 | >10 | >10 |
| P70S6K1 | 0.5 | C06 | -7 | -3 | -5 | 0.00159 | >10 | >10 |
| PKC-GAMMA | 0.5 | C06 | -7 | -3 | -5 | <0.000128 | >10 | >10 |
| CAMKK1 | 0.5 | C06 | -2 | -8 | -5 | 0.00465 | >10 | >10 |
| ALK4 | 0.5 | C06 | 2 | -13 | -6 | >10 | >10 | >10 |
| NEK6 | 0.5 | C06 | -2 | -9 | -6 | >10 | >10 | >10 |
| DAPK1 | 0.5 | C06 | -7 | -4 | -6 | 0.00260 | >10 | 7.31074 |
| HIPK4 | 0.5 | C06 | -4 | -7 | -6 | 0.22527 | >10 | 1.30449 |
| TLK1 | 0.5 | C06 | -8 | -4 | -6 | 0.02480 | >10 | >10 |
| EIF2AK4 | 0.5 | C06 | -9 | -3 | -6 | 0.23459 | >10 | >10 |
| PIM2 | 0.5 | C06 | -7 | -5 | -6 | 0.01671 | >10 | 1.22732 |
| CDK9-CYCLINT1 | 0.5 | C06 | -8 | -4 | -6 | 0.00521 | >10 | >10 |
| CAMK2G | 0.5 | C06 | -5 | -6 | -6 | <0.000128 | >10 | >10 |
| MAPKAPK-2 | 0.5 | C06 | -3 | -9 | -6 | 0.04190 | >10 | >10 |
| CK1 | 0.5 | C06 | -6 | -5 | -6 | 9.18203 | 1.08448 | >10 |
| GAK | 0.5 | C06 | -13 | 1 | -6 | 0.00538 | 0.07231 | 4.17885 |
| IRE1 | 0.5 | C06 | -7 | -5 | -6 | 0.09769 | >10 | >10 |
| PASK | 0.5 | C06 | -3 | -10 | -6 | 0.01843 | >10 | >10 |
| BLK | 0.5 | C06 | -18 | 6 | -6 | 0.00235 | >10 | >10 |
| NEK2 | 0.5 | C06 | -6 | -6 | -6 | 0.82949 | >10 | >10 |
| ICK | 0.5 | C06 | -5 | -7 | -6 | 0.00581 | >10 | >10 |
| HIPK2 | 0.5 | C06 | -9 | -4 | -7 | 0.41543 | >10 | 0.04962 |
| GSK-3-ALPHA | 0.5 | C06 | -6 | -8 | -7 | 0.00915 | >10 | 2.51914 |
| TSSK1 | 0.5 | C06 | -9 | -4 | -7 | <0.000128 | >10 | >10 |
| CLK4 | 0.5 | C06 | -14 | 0 | -7 | 0.00228 | >10 | 0.51890 |
| PKC-EPSILON | 0.5 | C06 | -10 | -3 | -7 | 0.00018 | >10 | >10 |
| MYO3A | 0.5 | C06 | -1 | -13 | -7 | 0.39527 | >10 | >10 |
| PDK2 | 0.5 | C06 | -4 | -10 | -7 | >10 | >10 | >10 |
| MRCKB | 0.5 | C06 | -10 | -4 | -7 | 0.00127 | >10 | >10 |
| ALK5 | 0.5 | C06 | 0 | -14 | -7 | 7.31590 | >10 | >10 |
| DYRK2 | 0.5 | C06 | 0 | -15 | -8 | 0.47149 | >10 | >10 |
| NDR1 | 0.5 | C06 | -8 | -8 | -8 | 0.00050 | >10 | >10 |
| DRAK1 | 0.5 | C06 | -7 | -8 | -8 | 0.01285 | >10 | >10 |
| CAMK4 | 0.5 | C06 | -9 | -7 | -8 | 0.28639 | >10 | >10 |
| SRPK1 | 0.5 | C06 | -6 | -10 | -8 | 0.72617 | >10 | >10 |
| CAMK2D | 0.5 | C06 | -11 | -4 | -8 | <0.000128 | >10 | >10 |
| MSK1 | 0.5 | C06 | -8 | -8 | -8 | 0.00034 | >10 | >10 |
| MARK3 | 0.5 | C06 | -3 | -14 | -8 | <0.000128 | >10 | >10 |
| PDK4 | 0.5 | C06 | -4 | -12 | -8 | >10 | >10 | >10 |
| MARK4 | 0.5 | C06 | -7 | -10 | -9 | <0.000128 | >10 | >10 |
| TAOK3 | 0.5 | C06 | -9 | -9 | -9 | 0.00320 | >10 | >10 |
| LATS1 | 0.5 | C06 | -3 | -14 | -9 | 0.00288 | >10 | >10 |
| CDK5-P35 | 0.5 | C06 | -4 | -13 | -9 | 0.00308 | >10 | >10 |
| NEK9 | 0.5 | C06 | -9 | -9 | -9 | 0.21784 | >10 | 0.46735 |
| PKC-IOTA | 0.5 | C06 | -3 | -15 | -9 | 0.01836 | >10 | >10 |
| PRKG2 | 0.5 | C06 | -9 | -9 | -9 | <0.000128 | 9.46865 | >10 |
| SGK3 | 0.5 | C06 | -11 | -7 | -9 | 0.15634 | >10 | >10 |
| MAP2K4 | 0.5 | C06 | -11 | -7 | -9 | 0.01019 | >10 | >10 |
| BTK | 0.5 | C06 | -16 | -2 | -9 | 0.11455 | >10 | >10 |
| TSSK3 | 0.5 | C06 | -11 | -8 | -9 | 0.00855 | >10 | >10 |
| NEK1 | 0.5 | C06 | -19 | 0 | -10 | 0.04597 | >10 | >10 |

| | | | | | | | | |
|---|---|---|---|---|---|---|---|---|
| P70S6K2 | 0.5 | C06 | -12 | -7 | -10 | 0.00246 | >10 | >10 |
| PLK4 | 0.5 | C06 | -15 | -4 | -10 | 0.00134 | >10 | >10 |
| PAK2 | 0.5 | C06 | -11 | -8 | -10 | 0.00341 | >10 | >10 |
| CK1-GAMMA3 | 0.5 | C06 | -9 | -11 | -10 | 5.19496 | >10 | 1.07631 |
| CRIK | 0.5 | C06 | -8 | -11 | -10 | 0.06940 | 1.00146 | >10 |
| ULK1 | 0.5 | C06 | -12 | -8 | -10 | 0.00698 | >10 | >10 |
| BUB1 | 0.5 | C06 | -12 | -8 | -10 | 0.50196 | 9.19314 | >10 |
| PAK5 | 0.5 | C06 | -12 | -8 | -10 | 0.00896 | >10 | >10 |
| NEK5 | 0.5 | C06 | -9 | -11 | -10 | 0.05773 | >10 | 1.75961 |
| GSK-3-BETA | 0.5 | C06 | 4 | -24 | -10 | 0.00555 | 5.78888 | 0.81312 |
| RSK1 | 0.5 | C06 | -8 | -12 | -10 | 0.00028 | >10 | >10 |
| BRSK2 | 0.5 | C06 | -10 | -11 | -10 | <0.000128 | >10 | >10 |
| PKC-ZETA | 0.5 | C06 | -9 | -12 | -11 | 0.10372 | >10 | >10 |
| DYRK3 | 0.5 | C06 | -11 | -11 | -11 | 0.01709 | >10 | >10 |
| PLK2 | 0.5 | C06 | -8 | -14 | -11 | 1.65830 | >10 | 0.08119 |
| CK1-GAMMA2 | 0.5 | C06 | -7 | -14 | -11 | 8.02594 | >10 | 1.35833 |
| PTK5 | 0.5 | C06 | -14 | -8 | -11 | 0.15230 | >10 | >10 |
| DAPK3 | 0.5 | C06 | -10 | -12 | -11 | 0.00264 | >10 | >10 |
| PRKG1 | 0.5 | C06 | -16 | -7 | -11 | 0.00104 | >10 | >10 |
| CLK3 | 0.5 | C06 | -11 | -12 | -11 | 3.17111 | >10 | 4.10123 |
| FER | 0.5 | C06 | -15 | -8 | -12 | 0.00510 | >10 | >10 |
| ROCK1 | 0.5 | C06 | -10 | -14 | -12 | 0.00072 | >10 | >10 |
| ALK2 | 0.5 | C06 | -11 | -12 | -12 | 1.94450 | >10 | >10 |
| IKK-EPSILON | 0.5 | C06 | -12 | -12 | -12 | 0.00091 | >10 | >10 |
| CDK3-CYCLINE | 0.5 | C06 | -12 | -11 | -12 | 0.00389 | >10 | >10 |
| PIM3 | 0.5 | C06 | -17 | -7 | -12 | <0.000128 | >10 | 2.29479 |
| TTBK2 | 0.5 | C06 | -14 | -10 | -12 | >10 | >10 | >10 |
| HIPK3 | 0.5 | C06 | -9 | -15 | -12 | 0.40118 | >10 | 0.05176 |
| INSR | 0.5 | C06 | -15 | -9 | -12 | 0.02076 | >10 | >10 |
| ALK6 | 0.5 | C06 | -15 | -9 | -12 | >10 | >10 | >10 |
| MSSK1 | 0.5 | C06 | -9 | -16 | -12 | >10 | >10 | >10 |
| LRRK2-G2019S | 0.5 | C06 | -17 | -8 | -12 | 0.00015 | >10 | >10 |
| STK25 | 0.5 | C06 | -15 | -10 | -12 | 0.00420 | >10 | >10 |
| mTORC1 | 0.5 | C06 | -8 | -17 | -12 | >10 | >10 | 1.86057 |
| IKK-BETA | 0.5 | C06 | -11 | -15 | -13 | 1.00921 | >10 | >10 |
| TBK1 | 0.5 | C06 | -24 | -1 | -13 | 0.00081 | >10 | >10 |
| QIK | 0.5 | C06 | -16 | -9 | -13 | 0.00075 | >10 | >10 |
| MINK | 0.5 | C06 | -12 | -14 | -13 | 0.00180 | >10 | >10 |
| DYRK4 | 0.5 | C06 | -11 | -15 | -13 | >10 | >10 | >10 |
| AMPK-A2B2G1 | 0.5 | C06 | -13 | -13 | -13 | <0.000128 | >10 | >10 |
| PKAC-BETA | 0.5 | C06 | -13 | -13 | -13 | 0.00093 | 3.35126 | >10 |
| IRAK4 | 0.5 | C06 | -7 | -19 | -13 | 0.00943 | >10 | >10 |
| AURORA-A | 0.5 | C06 | -14 | -12 | -13 | 0.00125 | >10 | >10 |
| PBK | 0.5 | C06 | -11 | -16 | -13 | 0.06670 | >10 | >10 |
| PLK3 | 0.5 | C06 | -9 | -18 | -13 | >10 | >10 | 0.15947 |
| DYRK1A | 0.5 | C06 | -13 | -13 | -13 | 0.00570 | >10 | >10 |
| PKN3 | 0.5 | C06 | -11 | -16 | -13 | 0.00015 | 0.06801 | >10 |
| CK1-EPSILON | 0.5 | C06 | -13 | -15 | -14 | 4.33692 | 1.22347 | >10 |
| GRK3 | 0.5 | C06 | -13 | -15 | -14 | 0.80100 | >10 | >10 |
| PDK3 | 0.5 | C06 | -14 | -14 | -14 | >10 | >10 | >10 |
| ROCK2 | 0.5 | C06 | -14 | -14 | -14 | 0.00032 | >10 | >10 |
| DCAMKL1 | 0.5 | C06 | -17 | -11 | -14 | 0.52181 | >10 | >10 |
| TESK1 | 0.5 | C06 | -13 | -15 | -14 | 0.26668 | >10 | >10 |
| NIM1K | 0.5 | C06 | -18 | -11 | -14 | 0.94607 | >10 | >10 |
| PI4-K-BETA | 0.5 | C06 | -15 | -14 | -15 | >10 | >10 | 0.05625 |
| IKK-ALPHA | 0.5 | C06 | -12 | -17 | -15 | 0.25709 | >10 | >10 |
| MYO3B | 0.5 | C06 | -16 | -14 | -15 | 0.16942 | >10 | >10 |
| NUAK2 | 0.5 | C06 | -12 | -18 | -15 | 0.00050 | >10 | >10 |
| AURORA-C | 0.5 | C06 | -13 | -17 | -15 | 0.00129 | >10 | >10 |
| HIPK1 | 0.5 | C06 | -12 | -18 | -15 | 0.93413 | >10 | 0.09903 |

| | | | | | | | | |
|---|---|---|---|---|---|---|---|---|
| MELK | 0.5 | C06 | -15 | -15 | -15 | <0.000128 | >10 | 2.64552 |
| PRKD2 | 0.5 | C06 | -12 | -20 | -16 | 0.00126 | >10 | >10 |
| CLK2 | 0.5 | C06 | -22 | -9 | -16 | 0.00527 | >10 | 0.84633 |
| AURORA-B | 0.5 | C06 | -21 | -10 | -16 | 0.00051 | >10 | >10 |
| PKAC-GAMMA | 0.5 | C06 | -22 | -10 | -16 | 0.00377 | 0.62978 | >10 |
| MLK3 | 0.5 | C06 | -16 | | -16 | 0.00843 | >10 | >10 |
| PI3K-BETA | 0.5 | C06 | -15 | -17 | -16 | >10 | >10 | 0.00041 |
| CK2A2 | 0.5 | C06 | -15 | -17 | -16 | 0.94840 | >10 | >10 |
| MST3 | 0.5 | C06 | -14 | -18 | -16 | 0.00207 | >10 | >10 |
| STK16 | 0.5 | C06 | -19 | -14 | -16 | 0.12130 | >10 | >10 |
| TSSK2 | 0.5 | C06 | -10 | -22 | -16 | 0.00386 | >10 | >10 |
| NDR2 | 0.5 | C06 | -15 | -18 | -16 | 0.00108 | >10 | >10 |
| PRAK | 0.5 | C06 | -17 | -16 | -16 | 0.52118 | >10 | >10 |
| PKN2 | 0.5 | C06 | -14 | -19 | -17 | 0.00021 | >10 | >10 |
| CAMK1D | 0.5 | C06 | -12 | -22 | -17 | 0.00318 | >10 | >10 |
| PAR1BA | 0.5 | C06 | -18 | -16 | -17 | <0.000128 | >10 | >10 |
| CDK6-CYCLIND3 | 0.5 | C06 | -11 | -23 | -17 | 0.03231 | >10 | >10 |
| PKA | 0.5 | C06 | -13 | -21 | -17 | 0.00047 | 3.62142 | >10 |
| SRPK2 | 0.5 | C06 | -18 | -16 | -17 | 1.08807 | >10 | >10 |
| PKC-ETA | 0.5 | C06 | -17 | -16 | -17 | <0.000128 | >10 | >10 |
| PKC-THETA | 0.5 | C06 | -16 | -18 | -17 | <0.000128 | >10 | 4.85311 |
| MEKK3 | 0.5 | C06 | -17 | -18 | -17 | 0.00780 | >10 | >10 |
| AMPK-A2B1G1 | 0.5 | C06 | -19 | -16 | -17 | <0.000128 | >10 | >10 |
| PKC-DELTA | 0.5 | C06 | -9 | -26 | -18 | <0.000128 | >10 | >10 |
| CDK2-CYCLINE | 0.5 | C06 | -17 | -18 | -18 | 0.00808 | >10 | >10 |
| CDK1-CYCLINB | 0.5 | C06 | -14 | -22 | -18 | 0.00346 | >10 | >10 |
| AMPK-A1B1G1 | 0.5 | C06 | -10 | -27 | -18 | <0.000128 | >10 | >10 |
| CAMK2A | 0.5 | C06 | -13 | -23 | -18 | <0.000128 | >10 | >10 |
| PHK-GAMMA1 | 0.5 | C06 | -19 | -18 | -18 | <0.000128 | >10 | >10 |
| PKN1 | 0.5 | C06 | -19 | -18 | -18 | 0.00013 | >10 | >10 |
| MAPK1 | 0.5 | C06 | -19 | -18 | -18 | 3.37047 | >10 | >10 |
| EIF2AK2 | 0.5 | C06 | -19 | -18 | -19 | 0.06776 | >10 | >10 |
| PHK-GAMMA2 | 0.5 | C06 | -20 | -17 | -19 | 0.00013 | >10 | >10 |
| CHEK2 | 0.5 | C06 | -21 | -16 | -19 | 0.01990 | >10 | >10 |
| CDK4-CYCLIND | 0.5 | C06 | -17 | -20 | -19 | 0.02734 | >10 | >10 |
| MLK2 | 0.5 | C06 | -18 | -20 | -19 | 0.00529 | 3.13215 | >10 |
| EIF2AK1 | 0.5 | C06 | -13 | -25 | -19 | 6.88377 | >10 | >10 |
| LATS2 | 0.5 | C06 | -14 | -24 | -19 | 0.00061 | >10 | >10 |
| CAMK1A | 0.5 | C06 | -14 | -24 | -19 | 0.07760 | >10 | >10 |
| CK1-GAMMA1 | 0.5 | C06 | -16 | -23 | -20 | >10 | >10 | 1.36284 |
| AKT3 | 0.5 | C06 | -17 | -23 | -20 | 0.00669 | >10 | >10 |
| SIK | 0.5 | C06 | -18 | -23 | -20 | 0.00035 | >10 | >10 |
| MAP4K2 | 0.5 | C06 | -19 | -21 | -20 | 0.00022 | >10 | >10 |
| PRKD1 | 0.5 | C06 | -18 | -24 | -21 | 0.00188 | >10 | >10 |
| MEK2 | 0.5 | C06 | -21 | -21 | -21 | 0.00042 | >10 | >10 |
| MAPK3 | 0.5 | C06 | -25 | -16 | -21 | 3.15115 | >10 | >10 |
| GRK7 | 0.5 | C06 | -29 | -13 | -21 | 0.00845 | >10 | >10 |
| PAK1 | 0.5 | C06 | -21 | -21 | -21 | 0.00141 | >10 | >10 |
| DGKz | 0.5 | C06 | -18 | -25 | -21 | >10 | >10 | >10 |
| ASK1 | 0.5 | C06 | -23 | -20 | -22 | 0.01159 | >10 | >10 |
| JNK2 | 0.5 | C06 | -21 | -23 | -22 | 6.19642 | >10 | >10 |
| PLK1 | 0.5 | C06 | -20 | -23 | -22 | 1.76491 | >10 | 0.03620 |
| MAPKAPK-3 | 0.5 | C06 | -26 | -18 | -22 | 3.26445 | >10 | >10 |
| JNK1 | 0.5 | C06 | -21 | -23 | -22 | 5.71654 | >10 | >10 |
| MAP3K8 | 0.5 | C06 | -31 | -13 | -22 | >10 | >10 | >10 |
| MST4 | 0.5 | C06 | -22 | -24 | -23 | 0.00324 | >10 | >10 |
| MEK1 | 0.5 | C06 | -24 | -22 | -23 | 0.00073 | >10 | >10 |

| | | | | | | | | |
|---|---|---|---|---|---|---|---|---|
| STK33 | 0.5 | C06 | -23 | -23 | -23 | 0.00703 | >10 | >10 |
| GRK5 | 0.5 | C06 | -21 | -27 | -24 | 0.03188 | >10 | >10 |
| NEK7 | 0.5 | C06 | -25 | -23 | -24 | >10 | >10 | >10 |
| DCAMKL2 | 0.5 | C06 | -26 | -23 | -25 | 0.00867 | >10 | >10 |
| PKC-BETA2 | 0.5 | C06 | -28 | -22 | -25 | 0.00053 | >10 | >10 |
| ARK5 | 0.5 | C06 | -19 | -32 | -25 | 0.00050 | >10 | >10 |
| PAK6 | 0.5 | C06 | -17 | -34 | -25 | 0.00249 | >10 | >10 |
| PRKX | 0.5 | C06 | -23 | -27 | -25 | 0.00076 | >10 | >10 |
| BMPR2 | 0.5 | C06 | -25 | -27 | -26 | >10 | >10 | >10 |
| JAK2 | 0.5 | C06 | -22 | -29 | -26 | 0.00015 | >10 | >10 |
| DRAK2 | 0.5 | C06 | -39 | -13 | -26 | 0.01250 | >10 | >10 |
| LIMK1 | 0.5 | C06 | -23 | -32 | -27 | 0.00243 | >10 | >10 |
| DLK | 0.5 | C06 | -21 | -33 | -27 | 1.83133 | >10 | >10 |
| CAMK2B | 0.5 | C06 | -28 | -27 | -27 | 0.00042 | >10 | >10 |
| MAP4K3 | 0.5 | C06 | -16 | -40 | -28 | <0.000128 | >10 | >10 |
| TTBK1 | 0.5 | C06 | -29 | -28 | -28 | >10 | >10 | >10 |
| MNK1 | 0.5 | C06 | -28 | -31 | -30 | 0.06993 | >10 | >10 |
| CHEK1 | 0.5 | C06 | -30 | -29 | -30 | <0.000128 | >10 | >10 |
| AMPK-A1B2G1 | 0.5 | C06 | -31 | -29 | -30 | <0.000128 | >10 | >10 |
| CDK2-CYCLINA | 0.5 | C06 | -31 | -29 | -30 | 0.00222 | >10 | >10 |
| JNK3 | 0.5 | C06 | -32 | -28 | -30 | 2.82612 | >10 | >10 |
| DGKa | 0.5 | C06 | -25 | -37 | -31 | >10 | >10 | >10 |
| mTOR | 0.5 | C06 | -40 | -26 | -33 | 1.10726 | >10 | 1.15961 |
| MARK1 | 0.5 | C06 | -32 | -34 | -33 | <0.000128 | >10 | >10 |
| PIKFYVE | 0.5 | C06 | -32 | -36 | -34 | 0.09161 | 0.01880 | 3.46847 |
| EIF2AK3 | 0.5 | C06 | -37 | -32 | -34 | >10 | >10 | >10 |
| GRK6 | 0.5 | C06 | -40 | -30 | -35 | 0.04276 | >10 | >10 |
| MLK1 | 0.5 | C06 | -46 | -28 | -37 | 0.00023 | 5.61304 | >10 |
| MEK3 | 0.5 | C06 | -39 | -38 | -38 | 0.05359 | >10 | >10 |
| WEE1 | 0.5 | C06 | -36 | -42 | -39 | 3.40721 | >10 | >10 |
| HASPIN | 0.5 | C06 | -41 | -42 | -41 | 0.01084 | >10 | >10 |
| EPH-A4 | 0.5 | C06 | -39 | -46 | -42 | 0.89313 | >10 | >10 |
| BRSK1 | 0.5 | C06 | -42 | -44 | -43 | 0.00036 | >10 | >10 |
| WNK1 | 0.5 | C06 | -42 | -49 | -45 | >10 | >10 | >10 |
| DGKb | 0.5 | C06 | -41 | -52 | -46 | >10 | >10 | >10 |
| CDK7 | 0.5 | C06 | -46 | -49 | -48 | 0.01884 | >10 | >10 |
| LKB1 | 0.5 | C06 | -34 | -69 | -52 | 0.00019 | >10 | >10 |

*empty well: knocked out data point

### Supplementary Table S3. In vitro absorption study

| Test Compound | MCULE I.D. | Test Concentration | Permeability ( $10^{-6}$ cm/s) | | | Percent Recovery(%) | | |
|---|---|---|---|---|---|---|---|---|
| | | | 1st | 2nd | Mean | 1st | 2nd | Mean |
| **A-B permeability (Caco-2, pH 6.5/7.4)** | | | | | | | | |
| D08 | MCULE-8132215280 | 1.0E-05 M | 0.16 | 0.17 | 0.2 | 74 | 89 | 82 |
| A01 | MCULE-6746692707 | 1.0E-05 M | 5.99 | 6.29 | 6.1 | 79 | 78 | 78 |
| C06 | MCULE-2965976181 | 1.0E-05 M | 0.50 | 0.53 | 0.5 | 24 | 25 | 25 |
| **B-A permeability (Caco-2, pH 6.5/7.4)** | | | | | | | | |
| D08 | MCULE-8132215280 | 1.0E-05 M | 0.45 | 0.50 | 0.5 | 111 | 103 | 107 |
| A01 | MCULE-6746692707 | 1.0E-05 M | 0.21 | 0.27 | 0.2 | 20 | 36 | 28 |
| C06 | MCULE-2965976181 | 1.0E-05 M | 5.38 | 4.52 | 4.9 | 17 | 20 | 19 |

The apparent permeability coefficent ($P_{app}$) of the test compound was calculated as follows:

$$P_{app}(cm/s)= \frac{V_R * C_{R,end}}{\Delta t} * \frac{1}{A * (C_{D,mid} - C_{R,mid})}$$

where $V_R$ is the volume of the receiver chamber. $C_{R,end}$ is the concentration of the test compound in the receiver chamber at the end time point, $\Delta t$ is the incubation time, and A is the surface area of the cell monolayer. $C_{D,mid}$ is the calculated mid-point concentration of the test compound in the donor side, which is the mean value of the donor concentration at time 0 minute and the donor concentration at the end time point. $C_{R,mid}$ is the mid-point concentration of the test compound in the receiver side, which is one half of the receiver concentration at the end time point. Concentrations of the test compound were expressed as peak areas of the test compound.

The recovery of the test compound was calculated as follows:

$$Recovery(\%)= \frac{V_D * C_{D,end} + V_R * C_{R,end}}{V_D * C_{D0}} * 100$$

where $V_D$ and $V_R$ are the volumes of the donor and receiver chambers, respectively. $C_{D,end}$ is the concentration of the test compound in the donor sample at the end time point. $C_{R,end}$ is the concentration of the test compound in the receiver sample at the end time point. $C_{D0}$ is the concentration of the test compound in the donor sample at time zero. Concentrations of the test compound are expressed as peak areas of the test compound.
